# Surface Engineering of FLT4-Targeted Nanocarriers Enhances Cell-Softening Glaucoma Therapy

**DOI:** 10.1101/2021.05.19.444878

**Authors:** Michael P. Vincent, Trevor Stack, Amir Vahabikashi, Guorong Li, Kristin M. Perkumas, Ruiyi Ren, Haiyan Gong, W. Daniel Stamer, Mark Johnson, Evan A. Scott

## Abstract

Primary open-angle glaucoma is associated with elevated intraocular pressure (IOP) that damages the optic nerve and leads to gradual vision loss. Several agents that reduce the stiffness of pressure-regulating Schlemm’s canal endothelial cells, in the conventional outflow pathway of the eye, lower IOP in glaucoma patients and are approved for clinical use. However, poor drug penetration and uncontrolled biodistribution limit their efficacy and produce local adverse effects. Compared to other ocular endothelia, FLT4/VEGFR3 is expressed at elevated levels by Schlemm’s canal endothelial cells and can be exploited for targeted drug delivery. Here, we validate FLT4 receptors as a clinically relevant target on Schlemm’s canal cells from glaucomatous human donors and engineer polymeric self-assembled nanocarriers displaying lipid-anchored targeting ligands that optimally engage this receptor. Targeting constructs were synthesized as lipid-PEGX-peptide, differing in the number of PEG spacer units (x), and were embedded in micelles. We present a novel proteolysis assay for quantifying ligand accessibility that we employ to design and optimize our FLT4-targeting strategy for glaucoma nanotherapy. Peptide accessibility to proteases correlated with receptor-mediated targeting enhancements. Increasing the accessibility of FLT4-binding peptides enhanced nanocarrier uptake by Schlemm’s canal cells while simultaneously decreasing uptake by off-target vascular endothelial cells. Using a paired longitudinal IOP study *in vivo*, we show this enhanced targeting of Schlemm’s canal cells translates to IOP reductions that are sustained for a significantly longer time as compared to controls. Histological analysis of murine anterior segment tissue confirmed nanocarrier localization to Schlemm’s canal within one hour after intracameral administration. This work demonstrates that steric effects between surface-displayed ligands and PEG coronas significantly impact targeting performance of synthetic nanocarriers across multiple biological scales. Minimizing the obstruction of modular targeting ligands by PEG measurably improved the efficacy of glaucoma nanotherapy and is an important consideration for engineering PEGylated nanocarriers for targeted drug delivery.

## INTRODUCTION

Glaucoma is the leading cause of irreversible blindness, lacks a cure, and is currently estimated to affect 76 million people worldwide^1^. The prevalence of glaucoma continues to rise, with the number of individuals living with the disease projected to increase to ~112 million people by 2040^1^. The hallmark of glaucoma is elevated intraocular pressure (IOP ≥ 21 mm Hg)^2^, which progressively damages retinal ganglion cells^3–6^. Multiple lines of evidence demonstrate mechanical dysfunction in pressure-regulating tissues contribute to this increase in IOP^7–16^. In the eye, IOP is determined by the balance between the rate of aqueous humor production by the ciliary body, and the rate of aqueous humor outflow through tissues of the conventional^17–20^ and uveoscleral^21–24^ outflow pathways. Ocular hypertension results from increased resistance to aqueous humor outflow through the conventional outflow pathway^25^ that is positioned at the iridocorneal angle of the anterior segment and consists of the trabecular meshwork, Schlemm’s canal, collector channels, intrascleral venous plexus and the aqueous veins. Decreased porosity^7–11^ of the Schlemm’s canal endothelium contributes to the increase in outflow resistance in glaucoma, and these porosity changes arise from an increase in Schlemm’s canal stiffness that impairs pore formation^15^. Most recently, it was demonstrated that the source of outflow resistance is localized to a region within 1 μm of the inner wall of the Schlemm’s canal endothelial surface^16^.

Currently, first and second line glaucoma therapeutics decrease IOP by either (i) decreasing aqueous humor inflow, or (ii) driving a greater proportion of the aqueous humor outflow through the unconventional outflow pathway^26^. However, these treatments do not address the source of the increased outflow resistance that contributes to ocular hypertension^26^. Cell softening agents, such as actin depolymerizers and rho kinase inhibitors, improve aqueous humor outflow^27–29^ by directly addressing dysfunction in the conventional outflow pathway. Cell softening agents are approved for clinical use in the US (Netarsudil^30–35^) and abroad (Ripasudil^36^), but suffer from side effects, including conjunctival hyperemia, conjunctival hemorrhage, cornea verticillata, eye pruritus, reduced visual acuity, and blurred vision^26^. Targeted drug delivery vehicles may address these issues by directing cell softening agents to their site of action while minimizing adverse events arising from off-target effects.

To target the delivery of cell softening agents to Schlemm’s canal cells, we previously developed a lipid-anchored targeting peptide^37^ that binds to FLT4/VEGFR3 receptors. FLT4 receptors are enriched on the Schlemm’s canal endothelium in animal models^38^. Most recently, FLT4 receptors have been detected in cultured Schlemm’s canal cells derived from healthy human donors^37^ and in human donor eyes by single cell RNA sequencing^39,40^. Despite this early work, multiple uncertainties remain regarding the use of a FLT4-targeting strategy. One uncertainty pertains to clinical utility and whether FLT4 receptors are expressed on the Schlemm’s canal surface in the eyes of glaucoma patients to permit the targeted delivery of cell softening drugs. Second, to maximize the efficacy and minimize the side effects of these agents, it is important that the engineered vehicles optimally deliver drug payloads to Schlemm’s canal cells instead of off-target endothelial cell types, in particular the corneal endothelium. Regarding our initial Schlemm’s canal-targeting approach, steric considerations involving the PEG chains of the drug delivery vehicle chassis and their potential for obstructing receptor access to the peptide ligand motivated us to rationally design alternative ligand structures to improve performance. Suboptimal receptor binding is a general problem with targeted drug delivery systems^41^, and it is often left unresolved due to a lack of methods for optimizing ligand interactions at the nanocarrier surface. While useful insights into the role of ligand length on organ-level nanocarrier accumulation are available in the literature^42^, empirical relationships between ligand steric effects at nanocarrier surfaces and their consequences on multiscale targeting performance and impact on clinically-relevant outcomes remains unestablished. Here, we address these issues by developing a novel assay to measure peptide accessibility to proteolysis, which enabled us to quantify the availability of ligands for binding events. We were particularly interested in determining whether lipid-anchored targeting constructs can be optimized to maximize binding to FLT4 receptors, and whether differences in receptor engagement produced measurable differences in drug delivery vehicle performance in a clinically relevant setting.

Using a multidisciplinary approach that included molecular characterization, *in vitro* targeting studies and direct evaluation in a murine model, we examined the potential for these optimized nanotherapeutics to target a cell softening agent to Schlemm’s canal *in vivo* and lower IOP. C57BL/6J mice were used as an animal model in these studies due to their similar conventional outflow pathway anatomy, physiology, and pharmacologic responses to humans^43–45^. This work holds important clinical implications for the use of cell softening nanotherapies as safe and efficacious tools for managing glaucoma. Furthermore, the rational design principles established herein, as well as the empirical evidence supporting their utility, can be harnessed to meet diverse challenges in drug delivery.

## EXPERIMENTAL SECTION

### Chemicals

Unless otherwise stated, all chemical reagents were purchased from the Sigma Aldrich Chemical Company.

### Solid phase peptide synthesis

Fmoc-N-amido-dPEG_6_-acid, Fmoc-N-amido-dPEG_24_-acid, and Fmoc-N-amido-dPEG_24_-amido-dPEG_24_-acid (Quanta Biodesign) were purchased for use in the synthesis of the PG6, PG24, and PG48 peptide constructs. Standard Fmoc solid phase peptide synthesis was performed to synthesize PG6, PG24, and PG24x2 (PG48) peptides on a 0.125 mmol scale. Information regarding each synthesized peptide is presented in **Table S2**. The chemical structure of each peptide is presented in **Figure S1**.

### Preparation of PEG-*b*-PPS micelle formulations

PEG_45_-*b*-PPS_23_ polymer was synthesized using established procedures^46,47^ (**Table S1**). Briefly, sodium methoxide was used to deprotect PEG thioacetate. This deprotected PEG thioacetate is then used to initiate the polymerization of propylene sulfide through an anionic ring opening polymerization reaction. MC nanocarriers were self-assembled from PEG_45_-*b*-PPS_23_ polymer^46,47^ via cosolvent evaporation using established protocols^48^. For uptake studies *in vitro* MCs were prepared to load DiI hydrophobic dye (Invitrogen), whereas formulations intended for IOP studies *in vivo* were prepared to co-load Latrunculin A (LatA; Cayman Chemical Company) and DiI. The formed MCs were split into aliquots of equal volume prior to the addition of peptide. PG6, PG24, or PG48 targeting peptides were dissolved in DMSO and were added to the specified MC aliquots at either a 1% or 5% molar ratio (peptide:polymer). Blank MC controls (lacking peptide) were included in cellular uptake studies. All formulations were prepared under sterile conditions and were filtered using a Sephadex LH-20 column.

### Determination of peptide concentration

Peptide concentration in purified MCs was determined by measuring tryptophan fluorescence (λ_Ex_ = 270 nm, λ_Em_ = 350 nm), calibrated against a peptide concentration series, using a SpectraMax M3 spectrophotometer. The peptide is readily detectable using these parameters (**Figure S3**). The concentration of peptide was determined using a simple linear regression model that was obtained by fitting the calibration data (see **Figure S4** for a representative calibration curve).

### Small angle x-ray scattering (SAXS)

SAXS was performed using synchrotron radiation at the DuPont-Northwestern-Dow Collaborative Access Team (DND-CAT) beamline at the Advanced Photon Source at Argonne National Laboratory (Argonne, IL, USA). A 7.5 m sample-to-detector distance, 10 keV (λ = 1.24 Å) collimated x-rays, and 3 s exposure time was used in all SAXS experiments. The *q*-range of 0.001-0.5 Å^-1^ was used to analyze scattering, and silver behenate diffraction patterns were used for calibration. The momentum transfer vector (q) is defined in **Equation 1**, θ denotes the scattering angle:

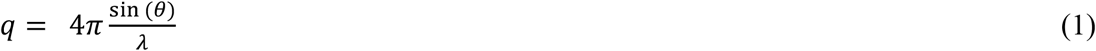

PRIMUS software (version 3.0.3) was used for data reduction. SasView (version 5.0) software was used for model fitting. A core shell sphere model (**Equation 2**) was fit to the scattering profiles of blank micelles (prepared without peptide), as well as micelles displaying PG6, PG24, or PG48 peptides at 1% or 5% molar ratios (peptide:polymer):

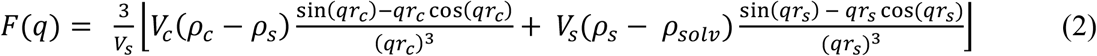

Where F is the structure factor, q is the momentum transfer vector (**Equation 1**), V_s_ is the particle volume (Å^3^), V_c_ is the particle core volume (Å^3^), ρ_c_ is the core scattering length density (10^-6^ Å^-2^), ρ_s_ is the shell scattering length density (10^-6^ Å^-2^), ρ_solv_ is the solvent scattering length density (10^-6^ Å^-2^), r_c_ is the core radius (Å). The radius of the total particle (r_s_; units: Å) is used to determine the shell thickness (r_t_; units: Å), as described by **Equation 3**:

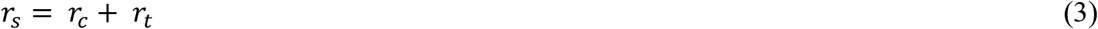

Micelle diameter values determined from DLS (**Table 2**) were used to select initial values for the core radius and shell thickness prior to parameter fitting. An iterative chi square (X^2^) minimization procedure using the Levenberg-Marquardt algorithm was used to fit optimal model parameters. A good core shell model fit is indicated by χ^2^ < 1.0. In all cases, χ^2^ << 0.1 was obtained for final fit models.

### Quantification of loaded drug concentration

Aliquots of purified LatA-loaded MC formulations were frozen at −80 °C and were lyophilized overnight. The resulting powder was resuspended in methanol and was placed at −20 °C for 1 h. Samples were centrifuged at 4,000 x g for 5 min to sediment the polymer. After this extraction procedure, the supernatant (containing drug) was collected for further analysis. The concentration of LatA was determined using high performance liquid chromatography (HPLC) calibrated against a concentration series of LatA prepared in methanol (**Figure S6**). The absorption of 235 nm light was measured. Data was acquired from three replicates. HPLC was performed using a C18 XDB-Eclipse column (Agilent) and a static methanol:water (95:5) mobile phase.

### Protease protection assay for examining the biochemical accessibility of targeting peptides

PEG-*b*-PPS micelle nanocarriers displaying PG6, PG24, or PG48 at 5% molar ratio were incubated with trypsin gold protease (Promega) at 37 °C, 80 rpm for 10 h. A peptide concentration of 40 nM and enzyme concentration of 800 nM (1:20 ratio) was used in these experiments. Reaction aliquots were quenched in 2% formic acid at the specified timepoints. Quenched reaction aliquots were mixed 1:1 with 50:50 methanol/acetonitrile with 0.1% trifloroacetic acid (TFA), and a saturating quantity of α-Cyano-4-hydroxycinnamic acid (Sigma) matrix. Samples were applied to 384-spot polished stainless-steel plates and were dried under hot air using a heat gun. Data was acquired using matrix assisted laser desorption ionization time-of-flight mass spectrometry (MALDI-TOF) using a Bruker rapifleX MALDI Tissuetyper TOF MS instrument.

### Primary cell culture

Normal and glaucomatous Schlemm’s canal cells were isolated from post-mortem human eyes obtained from BioSight (**Table 1**) and were cultured using established procedures^49,50^. Human umbilical vein endothelial cells (HUVECs) from pooled donors were purchased from Lonza, Ltd. The passage number of all primary cells used in these studies was less than six. Schlemm’s canal cells were cultured in low glucose DMEM (Gibco) supplemented with 10% fetal bovine serum and 1x penicillin-streptomycin-glutamine (Gibco). HUVECs were cultured in endothelial cell growth basal medium-2 (EBM-2; Lonza) supplemented with FBS and an EGM-2 BulletKit (Lonza) optimized for HUVEC culture. All cells were cultured at 37°C, 5% CO_2_ in T25 or T75 flasks.

**Table 1.** Donated human Schlemm’s canal (SC) endothelial cell lines used in this study.

| Cell Line | Disease State | Age | Gender<sup>†</sup> | Race<sup>†</sup> | Dex Test<sup>††</sup> | Eye Source | Eye Bank |
| --- | --- | --- | --- | --- | --- | --- | --- |
| SC 78 | Normal | 77 | Male | White | Negative | Whole eye | Kansas Eye Bank |
| SC 84 | Normal | 67 | NA | NA | Negative | Cornea | NC Eye Bank |
| SC 57g | Glaucoma | 78 | Male | NA | Negative | Whole eye | Life Legacy |
| SC 90g | Glaucoma | 71 | Female | White | Negative | Whole eye | NC Eye Bank |

<sup>†</sup>NA = Information not available
<sup>††</sup>Dex (dexamethasone) test. A negative Dex Test indicates a lack of myocilin induction upon treatment with dexamethasone, which is a response that is expected of SC cells and is used to distinguish isolated SC cells from the major possible contaminant, trabecular meshwork (TM) cells (a positive Dex Test is expected for TM cells).

### Quantification of FLT4/VEGFR3 expression by flow cytometry

Schlemm’s canal (SC) cells, glaucomatous Schlemm’s canal cells (SCg), or HUVECs (n=3) were seeded in 24-well plates at a density of 100,000 cells per well. Cells were cultured in their appropriate media types (described elsewhere in this methods section) and were allowed to adhere overnight at 37 °C, 5% CO_2_. On the following day, media was aspirated, cells were washed with PBS, and single cell suspensions were prepared following established procedures^37,51^. Cell pellets were resuspended in cell staining buffer with zombie aqua cell viability stain (BioLegend), were incubated at 4 °C for 15 minutes, and were washed with cell staining buffer. After a subsequent blocking step, cells were incubated with an APC-conjugated anti-human VEGFR3 (FLT4) antibody (BioLegend) for 20 minutes at 4 °C and were then subjected to two iterative rounds of washing (and centrifugation) per manufacturer recommendations. Cells were fixed using a paraformaldehyde cell-fixation buffer (BioLegend). Flow cytometry was performed using a BD LSRFortessa Flow Cytometer. Cytobank software^52^ was used to analyze the acquired data.

### Live cell imaging by high-throughput widefield fluorescence microscopy

Schlemm’s canal (SC) cells, glaucomatous Schlemm’s canal (SCg) cells, or HUVECs (n=3) were seeded in glass-bottom 96-well plates (Greiner) at a seeding density of 25,000 cells/well, and were placed at 37 °C, 5% CO_2_ overnight. Cells were washed, blocked, and were then placed in media containing APC-conjugated anti-human FLT4/VEGFR3 antibody (BioLegend) for 30 minutes at 37 °C. Afterwards, the media was removed by aspiration and cells were gently washed twice with fresh media. After the final wash step, cells were treated with media supplemented with cell-permeant NucBlue (Hoechst 33342) counterstain to visualize cell nuclei. High-throughput widefield fluorescence microscopy was performed using an ImageXpress High Content Imager (Molecular Devices). Images of live cells were acquired at 40X magnification in brightfield, DAPI, and Cy5 channels.

### Nanocarrier uptake studies

Schlemm’s canal cells or HUVECs were seeded at a density of 100,000 cells/well in 48-well plates and were allowed to adhere overnight at 37 °C, 5% CO_2_. Cells were treated with the specified DiI-loaded MC formulations (0.5 mg/mL polymer) for 2 h at 37 °C, 5% CO_2_. All experiments included untreated cells and a PBS-treatment group, as well as three biological replicates per treatment group (n=3). MC uptake was quantified by flow cytometry using a BD LSRFortessa Flow Cytometer and the acquired data was analyzed using Cytobank software^52^. The median fluorescence intensity (MFI) above PBS-treated background was calculated to remove cellular autofluorescence contributions to the measured values.

### Longitudinal evaluation of intraocular pressure (IOP) *in vivo*

The mice were anesthetized with ketamine (60 mg/kg) and xylazine (6 mg/kg). IOP was measured using rebound tonometry (TonoLab; Icare) immediately upon cessation of movement (i.e., in light sleep). Each recorded IOP was the average of six measurements, giving a total of 36 rebounds from the same eye per recorded IOP value. IOP was measured three times prior to nanocarrier treatment and 24, 30, 48, 72, and 96 h following nanoparticle injection. A total of nine mice were used for the 0-48 h timepoints. Four mice and three mice were carried through the 72 h and 96 h timepoints, respectively.

### Intracameral injection of nanocarrier formulations

For IOP studies, three-month-old female C57 mice were anesthetized with an intraperitoneal injection of ketamine (100 mg/kg) and xylazine (10 mg/kg). A drop of 0.5% proparacaine, a topical anesthetic, was applied to both eyes. Two pulled microglass needles filled with nanocarrier formulation (labeled “A” or “B”) and connected to a pump were alternatively inserted into both mouse anterior chambers. A volume of 2 μl of nanocarrier formulation “A” or “B” was infused into the anterior chamber of contralateral mouse eyes at a rate of 0.67 μl/min. After infusion, the needles were withdrawn, and topical erythromycin antibiotic ointment was applied to both eyes. All the mice were maintained on a warm water-circulating blanket until they had recovered from the anesthesia and the animals were subsequently returned to the animal housing rack. Unmasking of treatment identity occurred after all IOP measurements were recorded.

### Confocal microscopy of ocular tissues

All mouse eyes were dissected into 8 radial wedges. Each wedge was immersed in Vectashield ® mounting media with DAPI (Vector Laboratories) in a glass-bottom dish and imaged along one of the two sagittal planes with Zeiss LSM 700 confocal microscope (Carl Zeiss). The uninjected contralateral eyes were used as negative control. Single images and tiles images were taken with 20x objective to capture either the TM and its closely surrounding structures or the entire sagittal plane, using the ZEN2010 operating software (Carl Zeiss).

### Statistical analysis

Statistical analyses were performed using Prism software (version 9.0.0; GraphPad Prism Software, LLC). Details of each statistical analysis is provided in the corresponding figure legends.

## RESULTS

### Target validation: glaucomatous Schlemm’s canal endothelial cells highly express FLT4/VEGFR3 receptors

Schlemm’s canal endothelium is somewhat unique in that it expresses both vascular and lymphatic characteristics^53^. FLT4 receptors are a lymphatic marker expressed by Schlemm’s canal cells^38^. Developing strategies to promote nanocarrier interactions with FLT4 receptors offers a potential strategy for increasing their uptake by Schlemm’s canal endothelial cells and not nearby vascular structures of the iris and ciliary body. While past studies detected FLT4 receptors on the surface of Schlemm’s canal cells derived from normal human patients^37^, it is unclear whether glaucoma alters the expression of these receptors. To determine whether FLT4 is a viable cell surface marker for targeting Schlemm’s canal cells in patients with glaucoma, FLT4 expression was examined in two normal and two glaucomatous Schlemm’s canal cell strains from human donors (**Figure 1a**; **Table 1**).

**Figure 1.**
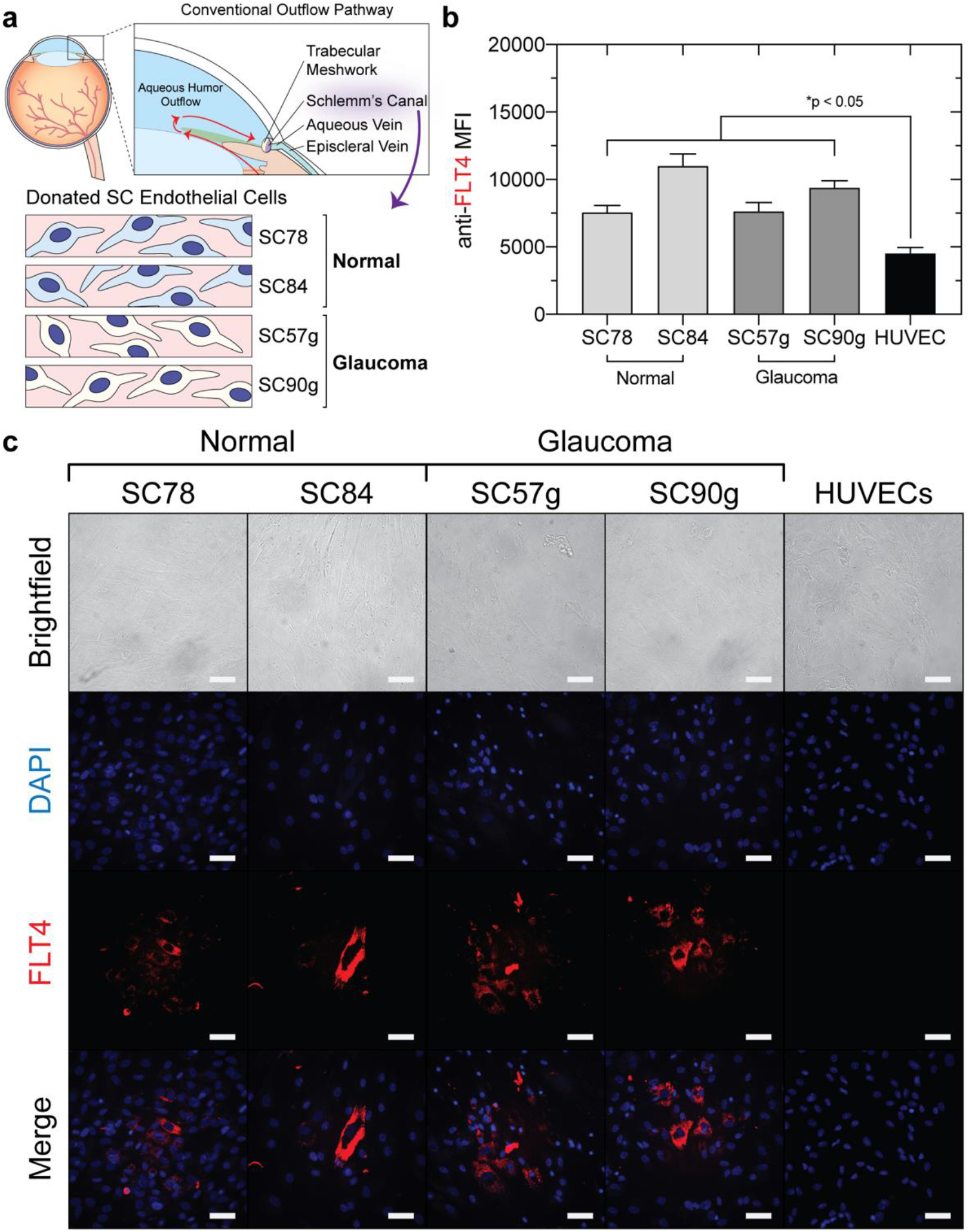
Normal and glaucomatous Schlemm’s canal (SC) endothelial cells express FLT4/VEGFR3. (**a**) Illustration of FLT4/VEGFR3 expression test in two normal SC endothelial cell strains (SC 78, SC 84) and two glaucomatous SC cell strains (SC 57g, SC 90g). (**b**) Flow cytometric analysis of FLT4 expression in SC and SC g cell strains. HUVECs are included as a control endothelial cell line that does not express FLT4 at high levels. FLT4 expression was detected by staining cells with antihuman FLT4-APC antibody. Data is presented as mean ± s.e.m. (n = 3). Significant differences between SC and HUVEC FLT4 MFI were determined by ANOVA with Dunnett’s multiple comparisons test (5% significance level). \**p*<0.05. (**c**) FLT4 expression observed by widefield fluorescence microscopy (40X). Brightfield images are shown together with the DAPI channel (cell nuclei) and Cy5 channel (anti-FLT4-APC). “Merge” denotes the merged DAPI and Cy5 channels. Scale bar = 50 μm.

Schlemm’s canal cells expressed FLT4 at significantly greater levels than human umbilical vein endothelial cells (HUVECs), a representative model of vascular endothelial cells, in both the normal and glaucomatous cell strains examined (**Figure 1b,c**). Quantification of FLT4 expression by flow cytometry (**Figure 1b**) was consistent with the FLT4 expression levels observed by widefield fluorescence microscopy (**Figure 1c**). While low FLT4 expression was detected on HUVECs by flow cytometry, this lower expression level is below the limit of detection of the less sensitive fluorescence microscopy technique. The relatively high FLT4 expression by Schlemm’s canal cells in the conventional outflow tissues^38^, and its maintenance during glaucoma (**Figure 1**), suggests FLT4 is available for targeting the delivery of IOP reducing agents directly to Schlemm’s canal cells in glaucoma patients.

### Development of Schlemm’s canal-targeted PEG-*b*-PPS micelles displaying optimized FLT4-binding peptides

The drug delivery vehicles developed in this work are polymeric micelles (MCs) self-assembled from oxidation-sensitive poly(ethylene glycol)-*b*-poly(propylene sulfide) (PEG_45_-*b*-PPS_23_) diblock copolymers (**Figure 2a-c**; **Table S1**). We hypothesized that the FLT4-binding peptides incorporating longer PEG spacers would more easily bind to the FLT4-binding receptors of the Schlemm’s canal cells as the longer spacer would minimize obstruction by the 45-unit micelle PEG corona. To test this hypothesis, peptide constructs were designed with 6- (PG6), 24- (PG24) or 48- (PG48) unit PEG spacers (**Figure 2d**; **Figure S1**). Peptides were synthesized using standard Fmoc solid phase peptide synthesis and the resulting products were of high purity (≥ 95% purity; **Figure 2d,e**). Dominant peaks of 2151.1, 3046.7, and 4071.4 Da are visible in the extracted mass spectra for PG6, PG24, and PG48, respectively, and these mass differences are consistent with the differences in the mass of the construct-specific PEG spacers (**Figure 2e**; **Table S2**).

**Figure 2.**
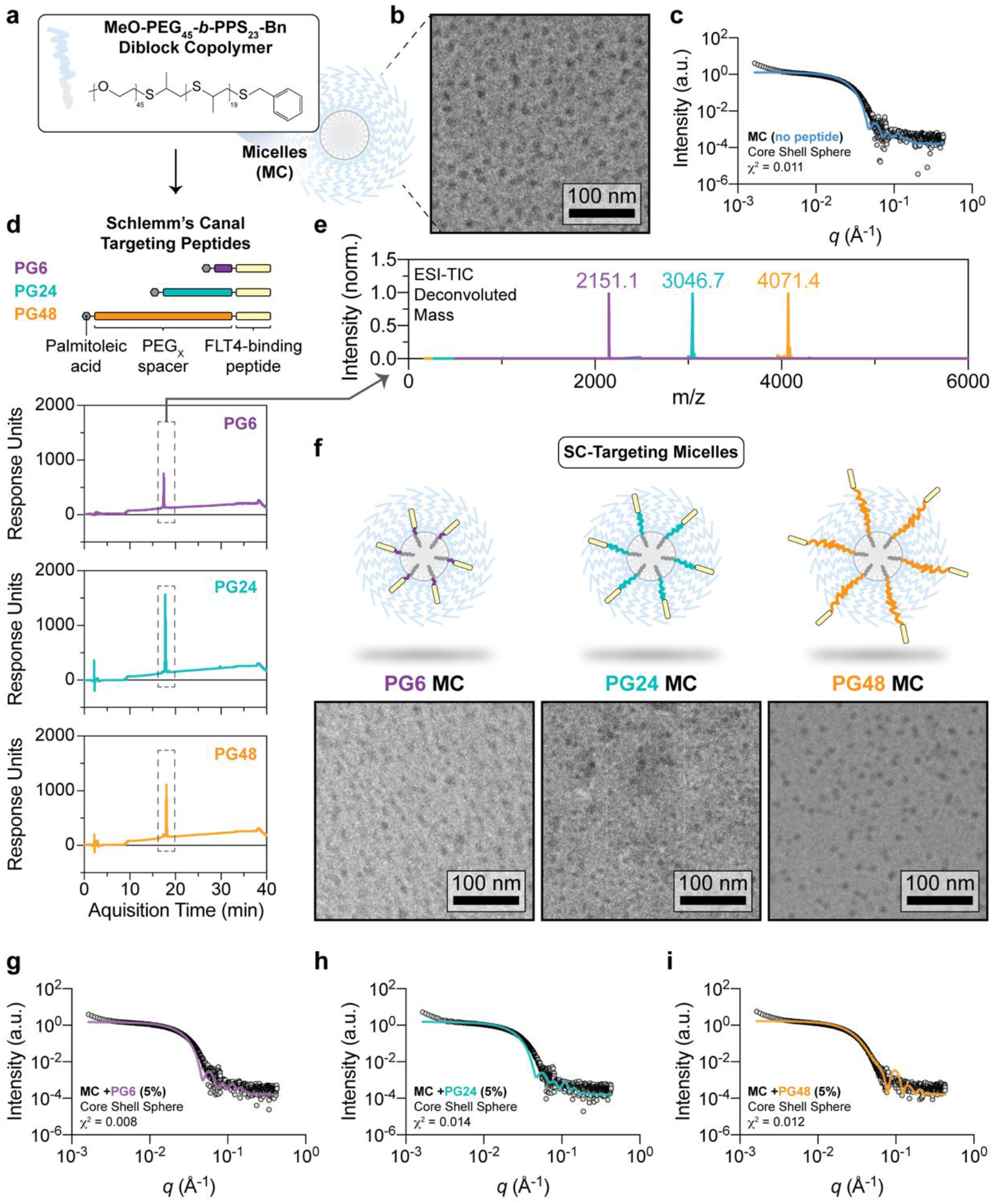
PEG-*b*-PPS micelles displaying lipid-anchored FLT4/VEGFR3-binding peptides that differ in the length of their PEG spacers. (**a-b**) Illustration of PEG-*b*-PPS micelles (**a**) and Cryo-TEM of blank micelles (i.e. without peptide) (**b**). (**c**) Micelle characterization by small angle x-ray scattering (SAXS). (**d**) Illustration of the designed FLT4-binding peptide constructs and LC-MS performed on the synthesized peptide products. (**e**) Deconvoluted mass spectra of purified peptides. Peaks are 2151.1, 3046.7, and 4071.4 Da for PG6, PG24, and PG48, respectively. (**f**) Cryo-TEM of MCs displaying the specified targeting peptides at 5% molar ratio (peptide:polymer). (**g-i**) Characterization of micelles displaying FLT4-binding peptides at a 5% molar ratio. The magnification is 10,000X and the scale bar is 100 nm for all Cryo-TEM micrographs presented. In all cases, small angle x-ray scattering (SAXS) was performed using synchrotron radiation at Argonne National Laboratory and a core shell model was fit to the data. The χ^2^ << 1.0 was obtained for all model fits (a good model fit is indicated by χ^2^ < 1.0).

Peptides were embedded into PEG_45_-*b*-PPS_23_ MC nanocarriers at 1% or 5% molar ratios (peptide:polymer) and were purified through a lipophilic Sephadex column to remove any unembedded peptide. The resulting micelle formulations were monodisperse (PDI < 0.1) with an average diameter of ~21-23 nm (**Table 2**). Electrophoretic light scattering (ELS) analysis demonstrate a zeta potential indicative of a neutrally charged surface for all nanocarriers in the presence and absence or peptide (**Table 2**). DLS and ELS procedures are described in the **Supplementary Methods**. Peptide incorporation into micelles was verified by Fourier transform infrared spectroscopy (FTIR; **Figure S2**; see **Supplementary Methods** for FTIR procedures) and peptide concentration was determined spectrophotometry (**Figure S3**; **Figure S4**). FTIR spectra of purified peptide-displaying micelle formulations revealed peaks in the amide I band (1700-1600 cm^-1^; C=O and C-N stretching vibrations) and amide II band (1590-1520 cm^-1^; N-H in-plane bending and C-N, C-C stretching vibrations) (**Figure S2**). These peaks are characteristic of the peptide bond and were absent from micelles prepared without peptide (**Figure S2**).

**Table 2.**
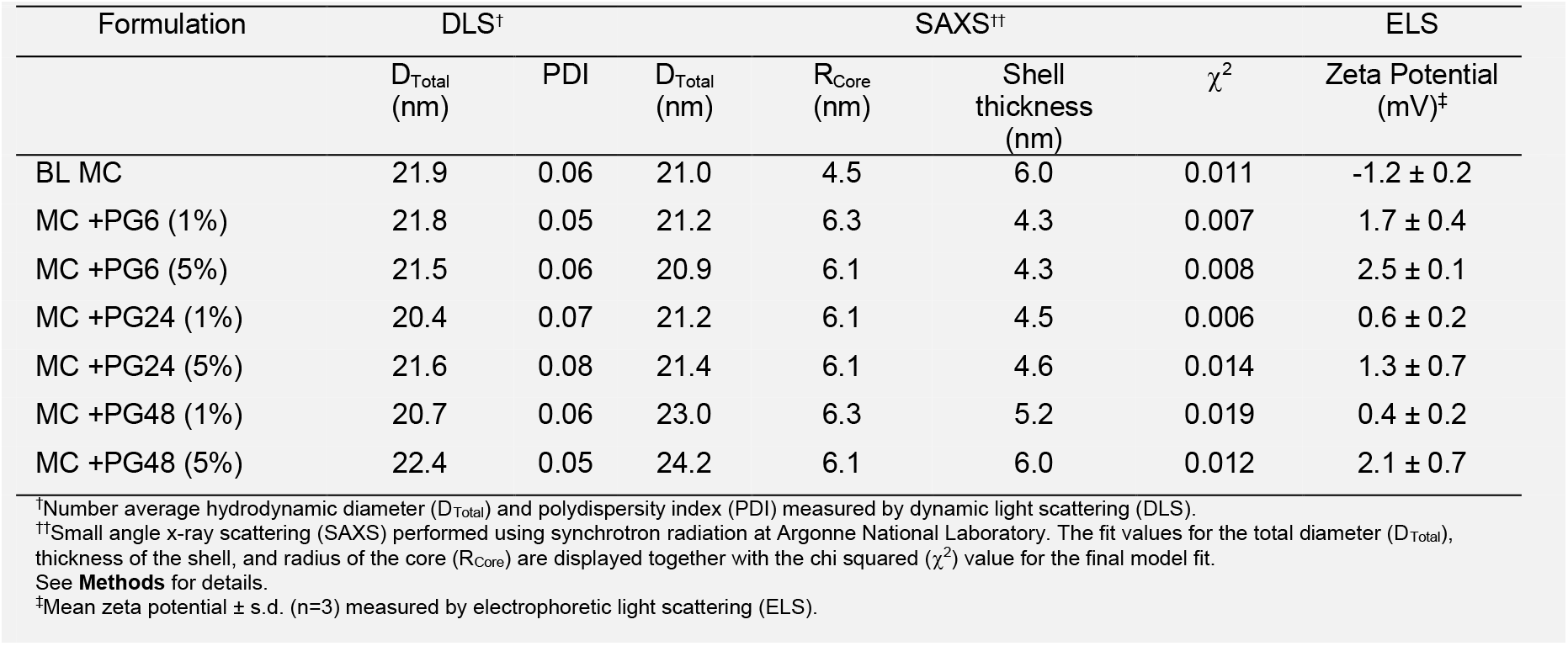
Physicochemical characterization of PEG-*b*-PPS micelles displaying FLT4-targeting peptide constructs.

The incorporation of peptide did not disrupt the spherical morphology expected for PEG-*b*-PPS micelles, as demonstrated by morphological analysis using cryogenic transmission electron microscopy (Cryo-TEM; **Figure 2f**; see **Supplementary Methods** for Cryo-TEM procedures) and small angle x-ray scattering (SAXS) performed using synchrotron radiation (**Figure 2g-i**; **Table 2**). In Cryo-TEM micrographs, PEG-*b*-PPS micelles appear as small dark circles in two-dimensions. These visible structures are the high contrast PPS hydrophobic core of the self-assembled PEG-*b*-PPS micelles, rather than the low contrast micelle PEG corona. Consistent with this interpretation, the hydrodynamic size measured by DLS (**Table 2**) exceeds the diameter of the PPS core observed by Cryo-TEM. The exclusive presence of the PPS core structures, and their size similarity across different formulations, is consistent with the expected micelle morphology. These results further suggest nanocarrier morphology is not perturbed by the palmitoleic lipid anchor of the peptide.

Micelle total diameter measurements obtained by SAXS was generally in agreement with the orthogonal analysis by DLS (**Table 2**). However, the higher resolution information obtained by probing the nanocarrier suspensions with high intensity x-rays did shed light on various structural details. Core shell models were fit to the data with high confidence, further supporting the micelle morphology was unperturbed by the presence of peptide at 1% (**Figure S5**) or 5% (**Figure 2g-i**) molar ratios. In all cases, the micelle core radius in the presence of peptide exceeded that of blank micelles by greater than 1 nm (**Table 2**). We attribute this observation to hydrophobic packing by the palmitoleic acid lipid anchor of the FLT4-binding peptides with the PPS core of the micelles.

While the shell thickness of micelles displaying PG48 at 1% or 5% molar ratio is greater than that of micelles displaying PG6 or PG24 at equivalent molar density, the shell thickness of PG48 formulations matched that of the blank micelles (**Table 2**). We attribute this observation to the greater continuity of the PEG corona in micelles prepared without peptide (blank micelles) and those displaying PG48, which scatter x-rays more consistently. The PG6 and PG24 constructs have fewer hydrophilic units than the 45-unit PEG block (PEG_45_) of the polymer (this is depicted in the **Figure 2f** illustrations). This length mismatch leaves a void volume that provides greater freedom for the PEG_45_ distal end to compact, leading to less consistent scattering near the micelle surface and a lower apparent shell thickness in these formulations. This interpretation will be corroborated by proteolysis experiments in the proceeding section, which probe the differences in biochemical access of each targeting ligand type.

### Evaluation of peptide biochemical accessibility and targeting Schlemm’s canal cells *in vitro*

We hypothesized that targeting peptides designed with longer PEG spacers would minimize ligand obstruction by the micelle PEG corona to biomolecules in the surrounding environment, and that these accessibility enhancements would improve peptide binding with FLT4 receptors (**Figure 1**). To examine differences in peptide accessibility, we developed a mass spectrometry-based proteolysis kinetic assay. This assay uses trypsin protease (23.3 kDa) as a biochemical probe (~5.8 nm diameter), which cleaves arginine (WHWLPNLRHYAS) in the peptide sequence to induce mass shifts that are detectable in quenched reaction aliquots (**Figure 3a**). A crude proteolysis rate is determined by monitoring the ratio of intact versus cleaved peptide signal intensities with time. Comparing differences in proteolysis kinetics provides a means to examine how accessible peptide constructs are to biomolecules in their environment when displayed on nanocarriers.

**Figure 3.**
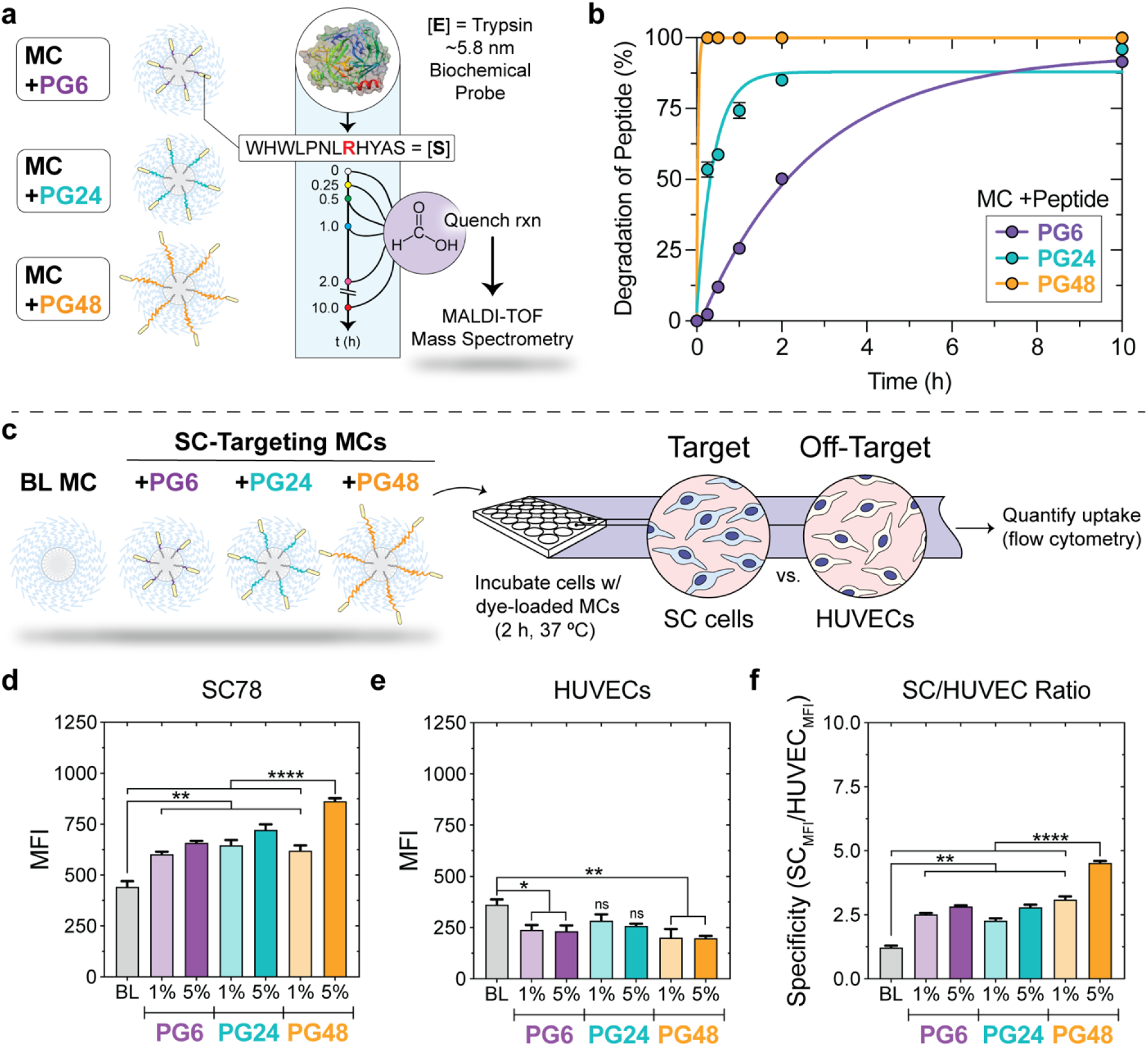
Differences in ligand biochemical accessibility modulates the rate of micelle uptake by Schlemm’s canal (SC) endothelial cells and vascular endothelial cells *in vitro*. (**a-b**) Determination of biochemical access to lipid-anchored targeting peptides displayed on polymeric micelles. (**a**) Illustration of protease protection assay to evaluate peptide accessibility. (**b**) Trypsin proteolysis kinetics (n=5). Concentrations: [Peptide] = 40 nM; [Trypsin] = 800 nM. Pseudo first-order association model fits are displayed for comparison, y = y_0_ + (y_max_-yo)*(1-e^-kx^), where k is the proteolysis rate (hours^-1^). In all cases, r^2^ > 0.94. (**c**-**f**) The surface-displayed PG48 FLT4-targeting peptide significantly increases micelle uptake by human SC cells and decreases uptake by HUVECs *in vitro*. (**c**) Illustration of PEG-*b*>-PPS micelle formulations and the cellular uptake study. (**d, e**) Cellular uptake by normal SC cells (**d**) or HUVECs (**e**). MFI determined by flow cytometry. The mean ± s.e.m. is displayed (n=3). (**f**) SC targeting specificity defined here as SC _MFI_/HUVEC_MFI_. For (**d-f**), statistical significance was determined by ANOVA with *post hoc* Tukey’s multiple comparisons test and a 5% significance level. ****p<0.0001, **p<0.01, *p<0.05.

When presented on micelles in the PG48 form, the FLT4-binding peptide substrate was cleaved to completion almost immediately, whereas buried PG6 and PG24 constructs were cleaved at much slower rates (**Figure 3b**). This result indicates protease access to the peptide component of the PG6 and PG24 constructs is obstructed by the 45-unit PEG corona of the MC nanocarriers to an extent that depends on the depth it is buried. The ligand obstruction observed for PG6 and PG24 constructs further suggests the potential for suboptimal engagement with FLT4/VEGFR3 receptors on the surface of the Schlemm’s canal cells.

To examine whether these differences in ligand accessibility translate into greater interactions with the Schlemm’s canal cells, we performed nanocarrier uptake studies with cultured Schlemm’s canal cells from normal human donors (target cell type) and HUVECs (vascular endothelial cell model; off-target cell type) (**Figure 3c**). Increasing peptide accessibility improved Schlemm’s canal -targeting *in vitro*. Compared to all other formulations, MC +PG48 (5%) significantly enhanced micelle uptake by Schlemm’s canal cells (**Figure 3d**) and decreased off-target uptake by HUVECs (Figure 3e). Micelles displaying the PG48 peptide resulted in a nearly 2-fold enhancement in Schlemm’s canal-targeting specificity compared to the PG6 and PG24 constructs that are buried within the micelle PEG corona (**Figure 3f**) and a 4.5-fold enhancement in Schlemm’s canal cell uptake over HUVEC (**Figure 3f**).

For micelles displaying the PG6 or PG24 constructs, the use of shorter PEG spacers together with the 12 amino acid FLT4-binding peptide obstructs biochemical access to the ligand (**Figure 3b**) and the extent of this obstruction is a function of ligand display depth within the micelle PEG corona. Importantly, differences in ligand biochemically accessibility resulted in significant differences in Schlemm’s canal cell targeting performance (**Figure 3d-f**). The targeting performance of a nanocarrier measurably decreased with increasing ligand obstruction.

### Performance evaluation of PG48 versus PG6 on enhancing the micellar delivery of IOP-reducing agents *in vivo*

As our goal was to develop nanocarriers capable of targeting Schlemm’s canal cells and delivering payloads that lower intraocular pressure, we sought to determine if the optimized nanocarriers displaying the FLT4-binding peptide above the surface would outperform our original, buried peptide construct (PG6) in a more clinically relevant setting. We were particularly interested in evaluating the performance of the original PG6 construct^37^ against the more highly-targeted PG48 construct in the delivery of the IOP-reducing agent latrunculin A (LatA) *in vivo*. LatA is a 16-membered macrolide that acts by blocking the incorporation of actin monomers into actin filaments and thereby depolymerizing actin^54–57^ and lowering cell stiffness^37,48^. We executed a longitudinal assessment of IOP in C57BL/6J mice receiving LatA MC displaying either PG6 or PG48 in contralateral eyes. Aside from the introduction of the optimized targeting construct, this study incorporated a paired design and greater statistical power than our previous analysis^37^. Animal information and maintenance procedures can be found in the **Supplementary Methods**.

The baseline IOP was measured by tonometry prior to treatment. Significant differences were not found in the contralateral IOP measured at baseline (OS: 19.9 ± 0.1 mmHg; OD: 19.9 ± 0.2 mmHg; **Figure S7**). Afterwards, LatA MC +PG48 and LatA MC +PG6 were administered intracamerally to contralateral eyes (i.e., one eye received one formulation type whereas the other eye received the second formulation type) (**Figure 4a**). IOP was measured by tonometry at 24, 30, 48, 72, and 96 h following intracameral injection (**Figure 4a**). Compared to baseline IOP values, both formulations led to a significant decrease in IOP at the 24 h timepoint, with an IOP reduction of −7.5 ± 0.9 mmHg in the LatA MC +PG48 treatment group and −7.1 ± 1.4 mmHg in the LatA MC +PG6 group (**Figure 4b**). While the decrease in IOP in the PG48 group was greater at 24 h than the PG6 group, this difference was not statistically significant.

**Figure 4.**
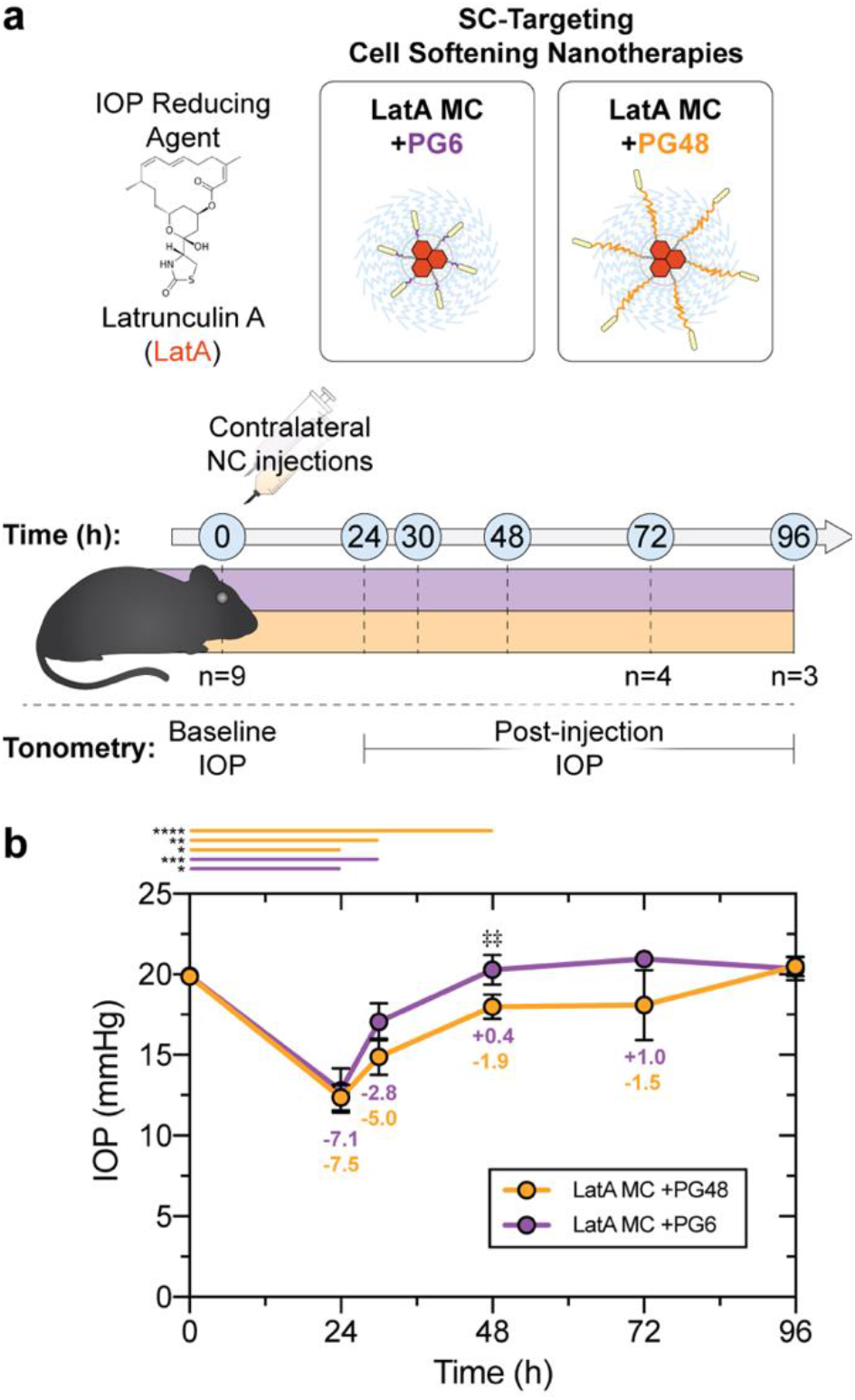
Increasing the biochemical accessibility of the FLT4-targeting peptide on SC-targeting nanocarriers enhances efficacy of a model IOP-reducing agent *in vivo*. (**a**) Experimental overview. The performance of LatA-loaded micelles (18 μM LatA) displaying either the PG6 or PG48 FLT4-binding peptide was evaluated *in vivo* in a paired IOP study. Nanocarriers were injected intracamerally into the contralateral eyes of mice. IOP was measured prior to injection (baseline), and at 24, 30, 48, 72, and 96 h after injection. Nine mice (n=9) were evaluated through 48 h. Measurements from four (n=4) and three (n=3) mice were obtained at 72 h and 96 h timepoints, respectively. (**b**) IOP time course at baseline and after intracameral injection of the specified nanocarrier formulations. Statistically significant differences between the PG6 and PG48 treatment groups was determined using a paired, twotailed t-test and a 5% significance level. ^‡‡^*p* < 0.01. The bars above the plot show statistically significant differences in IOP from the baseline value within the specified treatment group, assessed using a paired, two-tailed t-test. \*\*\*\**p* < 0.0001; \*\*\**p* ≤ 0.001; \*\**p* < 0.005; \**p* < 0.05.

However, a notable divergence in the efficacy of the two competing formulations was observed after 24 h (**Figure 4b**). LatA-loaded micelles displaying the PG48 peptide sustained a reduction in IOP through 72 h and did not return to baseline until day 4 post-treatment (**Figure 4b**), in contrast to the PG6 treatment group, in which the IOP returned to baseline by 48 h (**Figure 4b**). Our paired statistical analysis demonstrated that the IOP of eyes treated with LatA MC +PG48 (18.0 ± 0.7 mmHg) was significantly lower than those treated with LatA MC +PG6 (20.3 ± 0.9 mmHg) at 48 h (**Figure 4b**). Furthermore, the IOP measured in eyes receiving LatA MC +PG48 was significantly lower than baseline values through 48 h (i.e., at 24, 30, and 48 h) (**Figure 4b**). For eyes receiving LatA MC +PG6, significant differences from baseline IOP were only observed through 30 h (**Figure 4b**).

From these results, we conclude the targeting performance of micelles displaying the PG48 FLT4-binding peptide is superior to that of micelles displaying the PG6 variant. This improved ability to target cell softening agent delivery to the Schlemm’s canal endothelium enhanced efficacy by achieving a prolonged reduction in IOP that was statistically significant through 48 h after a single injection.

### Localization of the optimal SC-targeting nanocarriers in the murine conventional outflow pathway

We next examined the localization of micelles displaying the PG48 peptide within the murine conventional outflow pathway (**Figure 5a**). We were particularly interested in the localization of PG48-displaying micelles at an early timepoint that would precede its observed effects in our longitudinal IOP assessment (**Figure 4**). To this end, PG48-displaying micelles were prepared to load a lipophilic dye to permit fluorescence tracing in conventional outflow tissues. Nanocarriers were administered intracamerally into C57BL/6J mice and eyes were enucleated at 45 minutes post-injection for histology (see **Supplementary Methods** for ocular histology procedures).

**Figure 5.**
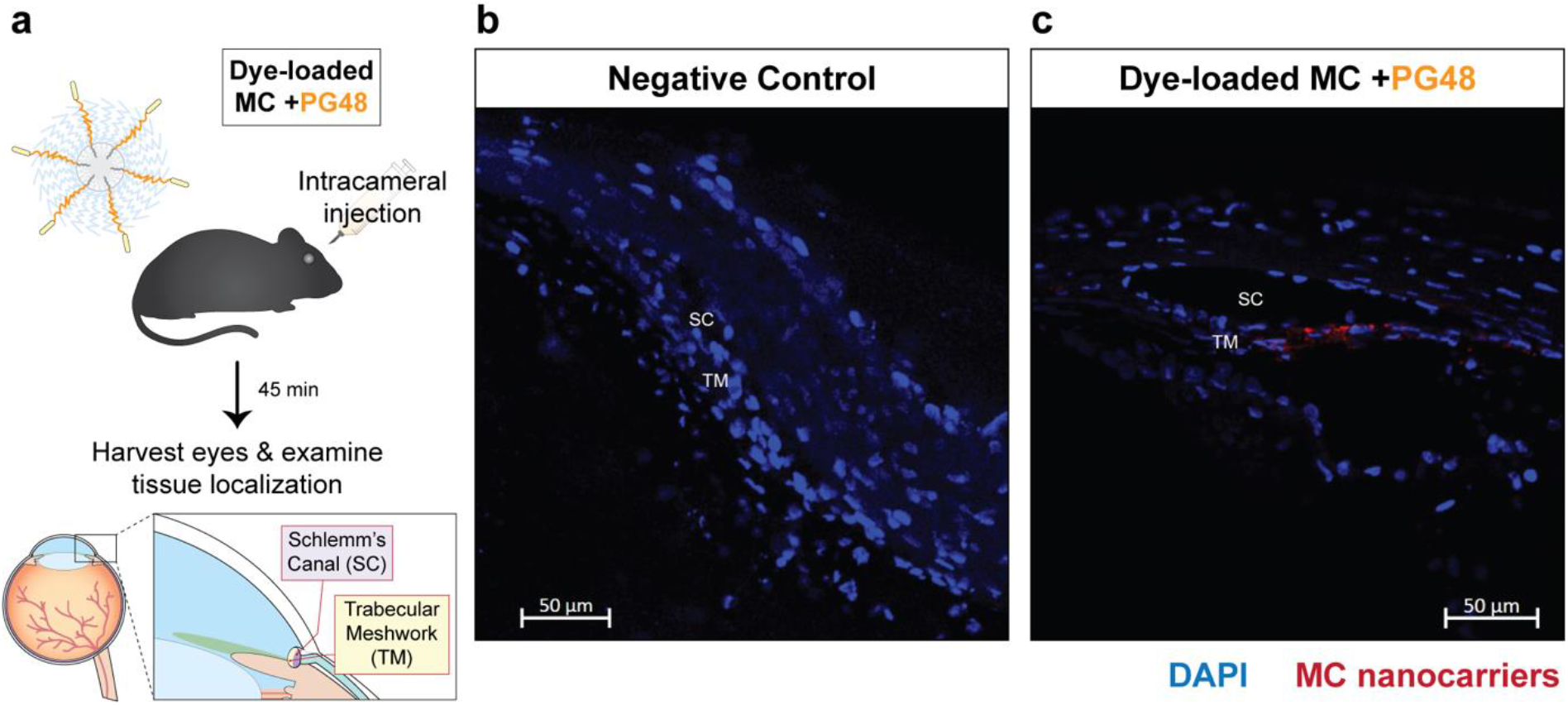
Histological analysis of MC +PG48 localization within the murine conventional outflow pathway *in vivo*. (**a**) Experimental illustration. DiI dye-loaded MC +PG48 (5%) nanocarriers (20 mg/mL polymer concentration) were administered intracamerally in C57BL/6J mice. After 45 min, eyes were enucleated and fixed for confocal microscopy analysis. (**b**, **c**) Confocal microscopy of conventional outflow pathway tissues obtained from eyes of (**b**) the negative control group (no injection) and (**c**) the MC +PG48 treatment group. Images were acquired at 20X magnification. DAPI and DiI signals are overlaid to present the cell nuclei and nanocarrier localization, respectively. Scale bar = 50 μm. Tissue abbreviations: SC: Schlemm’s canal; TM: Trabecular meshwork.

We examined tissue sections prepared from the anterior segments of the negative control and nanocarrier-receiving eyes (**Figure 5b,c**). The PG48-displaying micelles were distributed non-uniformly circumferentially around the eye (data not shown). These SC-targeting nanocarriers accumulated at the inner wall of Schlemm’s canal and the neighboring juxtacanalicular tissue (JCT) of the trabecular meshwork (TM) (**Figure 5c**). Interestingly, the fluorescent signal that localized to the SC was much greater than the TM-localizing signal (**Figure 5c**). We further note that at this timepoint (45 min post-injection), the nanocarriers were also observed to a lesser extent along the endothelial side of cornea but were blocked by the endothelial barrier. From this analysis, we conclude that the SC-targeting micelles accumulate at the SC endothelium at higher concentrations than in the nearby trabecular meshwork. The relatively strong localization to the SC in the anterior chamber further suggests that the FLT4-binding strategy achieves high specificity for the tissue of interest.

## DISCUSSION

Glaucoma is a progressive disease that, if untreated, slowly but relentlessly destroys ganglion cells and robs vision. The lowering of IOP is the only treatment for glaucoma that is proven to decrease disease progression. Agents such as actin depolymerizers and Rho kinase inhibitors reduce cell stiffness and have been shown to lower the IOP when delivered to the aqueous humor outflow pathway. However, these agents have significant side effects that are associated with off-target vasodilation of ocular surface vessels. Using optimized nanocarriers loaded with an actin depolymerizing agent, LatA, we demonstrate here that we can target Schlemm’s canal cells in a highly specific fashion using versatile nanocarriers and that these nanocarriers when loaded with a cell softening agent significantly lower IOP in murine eyes for an extended period of time.

As the goal was to develop a nanocarrier that selectively targeted Schlemm’s canal cells, it was necessary to demonstrate that the target, FLT4, is expressed at high levels not only on normal, but also glaucomatous Schlemm’s canal cells (**Figure 1**). This indicated that promoting nanocarrier engagement with these receptors may provide a clinically useful approach to direct the delivery of cell softening agents to the Schlemm’s canal cells. These data also suggested that the disease state does not hinder the expression of FLT4 by Schlemm’s canal cells.

The overarching goal of the present work was to develop an improved nanotechnology platform for targeting the delivery of cell softening agents to the Schlemm’s canal endothelium in order to lower IOP. Previous studies of LatA demonstrated promise in reducing the IOP ^28,29^ but did not lead to the development of a clinical product. Our previous biomechanical analyses using atomic force microscopy (AFM) demonstrated that untargeted micellar delivery of LatA softens Schlemm’s canal cells *in vitro*^48^, thereby confirming the successful lysosomal escape of an actin depolymerizing agent following cellular internalization and validated PEG-*b*-PPS nanocarriers as appropriate vehicles for delivering cell softening agents. In a separate set of studies we demonstrated that the targeted micellar delivery of LatA (LatA MC +PG6) at a low concentration of 50 nM is sufficient to significantly reduce Schlemm’s canal cell stiffness compared to control *in vitro*^37^, and the targeted micellar delivery of 15.5-17 μM LatA using PG6 led to significant reductions in murine IOP for up to 24 h *in vivo* following intracameral injection^37^.

To improve the efficacy of cell softening agents delivered by Schlemm’s canal-targeting micelles, we rationally designed variants of our modular lipid-anchored FLT4-binding peptide to achieve a greater influx of drug-loaded nanocarriers into the Schlemm’s canal cells. We began this design process with the observation that our original Schlemm’s canal-targeting construct^37^, denoted in the present work as “PG6”, was likely buried beneath the micelle surface where it is less accessible to FLT4 receptors. This design consideration is fairly unique to lipid-anchored targeting peptides embedded in amphiphilic nanocarriers, since materials are more commonly prepared for targeting via the direct conjugation of ligand to the surface^58^—a strategy that can encounter issues with a lack of modularity, usage of harsh chemical conditions, control over display density, among others^59^.

Understanding the relationship between the biochemical access of a lipid-anchored ligand and targeting performance became a focus of the present work, since the use of targeting moieties to direct vehicles to the intended location requires (i) successful ligand docking with the cognate receptor, which is enabled by the specific arrangement of functional groups of the ligand with those of the receptor binding pocket, but also (ii) a lack of obstruction that hinders formation of the ligand-receptor complex. The latter consideration permits successful binding events to occur with a greater frequency. Whereas the first requirement is fulfilled by the WHWLPNLRHYAS peptide that was developed by phage display to specifically bind FLT4 receptors, the second requirement was not fully met, since the use of the six-unit PEG spacer left the lipid-anchored peptide shrouded within the 45-unit PEG corona of the micelles self-assembling from PEG_45_-*b*-PPS_19_ polymer.

These considerations prompted our rational design of optimized targeting peptide constructs that differed in their PEG linker lengths and display depth on micelles (**Figure 2**). As a pre-requisite for understanding the influence of these steric differences on targeting performance, we required an assay that is capable of probing peptide accessibility at the nanoscale. We sought to develop an assay that is high-throughput compatible and avoids the use of receptor proteins, which are notoriously difficult to purify in their native/active form and are expensive when commercially available. We reasoned the accessibility of nanocarrier-displayed peptide to its cognate receptor would correlate with its susceptibility to proteolysis *in vitro*. Despite being very different in nature, receptor binding and protease-mediated cleavage both require physical access to the peptide ligand to initiate a successful interaction. It seemed plausible to quantify peptide accessibility by monitoring the occurrence of a peptide-modifying event with time. If the steric differences influenced peptide access, the frequency of a peptide-modifying event should decrease with greater obstruction by the micelle PEG corona.

These considerations inspired our development of a novel method that uses trypsin as a biochemical probe. Trypsin acts on each peptide molecule only once, since the peptide contains one arginine residue and lysine is absent. Cleavage of the peptide by trypsin produces irreversible mass shifts, and the proteolysis kinetics are monitored to quantify differences in the accessibility of lipid-anchored targeting peptides displayed on amphiphilic nanocarriers (**Figure 3a**). In this assay, we hypothesized that PG48 is displayed at the micelle surface where it is most susceptible to proteolysis, thereby yielding peptide cleavage at a greater rate than the PG24 or PG6 peptides. PG24 and PG6 incorporate shorter PEG linkers (**Figure 2**), leaving these constructs buried to different extents within the outer PEG corona micelles that self-assemble from PEG_45_-*b*-PPS_19_.

In agreement with our hypothesis, the protease protection kinetic assay developed herein demonstrated that the use of 6-, 24-, or 48-unit PEG spacers achieves significant differences in biomolecular access to the peptide ligand (**Figure 3a,b**). These molecular-level differences in ligand access translated to differences in targeting specificity *in vitro* in studies conducted with primary cultures of Schlemm’s canal cells obtained from human donors (**Figure 3c-f**). Micelles displaying the most accessible targeting peptide, PG48, achieved the greatest targeting specificity for Schlemm’s canal cells in these studies (**Figure 3f**). This specificity improvement resulted in both an increase in uptake by the Schlemm’s canal cells (**Figure 3d**) and a decrease in uptake by vascular endothelial cells, a model of cells where uptake is not desirable^26^ (**Figure 3e**).

Our paired longitudinal IOP studies demonstrated that LatA-loaded micelles displaying the biochemically enhanced PG48 construct led to a 7.5 mmHg reduction in IOP, on average, after 24 h compared to 7.1 mmHg achieved by treatment with the LatA MC+PG6 formulation (**Figure 4b**). While this change in IOP was not significantly different between the two groups at 24 h, the greater magnitude decrease observed after treatment with LatA MC+PG48 is consistent with a greater amount of drug reaching Schlemm’s canal endothelium *in vivo*. Strikingly, mice receiving the optimized drug delivery vehicle maintained IOP reductions of −5.0, −1.9, and −1.5 mmHg at 30, 48, and 72 hours, respectively, with the return to baseline observed at 96 hours post-administration (**Figure 4b**). The IOP reductions at 24, 30, and 48 hours differed significantly from baseline levels for the PG48 group. In contrast, contralateral eyes receiving LatA MC+PG6 returned to baseline by 48 h after achieving a significant decrease in IOP at 24 h and 30 h (**Figure 4b**)—an outcome that is consistent with the transient IOP decrease observed in our past reports from two separate trials with this construct^37^. Importantly, the IOP reduction in the PG48 group was significantly greater in magnitude than the PG6 group at the 48-hour timepoint (**Figure 4b**). Collectively, these results demonstrate the superiority of the PG48 construct for targeting the micellar delivery of a cell softening glaucoma therapeutic to Schlemm’s canal in the eye. Histological analysis confirmed that micelles displaying the PG48 FLT4-binding peptide accumulate within Schlemm’s canal *in vivo* (**Figure 5c**).

Enhancing the biochemical access to the FLT4 binding peptide leads to a greater accumulation of cell softening agents within the Schlemm’s canal endothelium. Higher drug concentrations at the target site resulted in longer lasting improvements in outflow facility to yield an IOP reduction that is sustained for a longer period of time. We further conclude that micelles displaying the PG48 construct are the highest performing nanocarrier chassis for delivering IOP-lowering cell softening agents to Schlemm’s canal. PEG-*b*-PPS micelles efficiently load diverse hydrophobic small molecules^47,60^ and will accommodate a wide variety of new and currently available therapeutics^61^.

Looking ahead, there remains the practical challenges of how the therapeutic effect of softening Schlemm’s canal cells can be sustained for a longer period of time. As the corneal endothelium is permeable only to molecules with molecular weights less than approximately 500 Daltons^62,63^, nanoparticles cannot be delivered to the anterior segment topically. Instead, these agents need to be either (i) injected to the anterior segment or retro-injected into the aqueous veins^64^, or (ii) released continuously from a long-term depot/device placed in the eye. While injection is plausible for targeted-nanocarriers carrying gene therapy to Schlemm’s canal, long term release is required for pharmacological treatments such as LatA carrying micelles as frequent intraocular injections would not be tolerated.

Aside from the more conventional delivery of pharmacological payloads, our Schlemm’s canal targeting strategy can be extended to the development of gene delivery technologies^65^. Glaucoma is an attractive target for gene therapy^66,67^ and modulating the expression of genes that regulate the stiffness of Schlemm’s canal endothelial cells may hold the key to producing more permanent enhancements in outflow facility without stents or surgery. The lipid-anchored FLT4-binding peptides are modular, and readily incorporate into alternative PEG-*b*-PPS morphologies, such as vesicular polymersomes^51,59,60,68,69^ or bicontinuous nanospheres^70–72^, which are capable of encapsulating the hydrophilic cargoes necessary for genetic intervention. This includes regulatory RNAs for the transient modulation of gene expression, or CRISPR/Cas9 components for stable genome editing.

## CONCLUSION

In closing, we demonstrate that steric effects between surface-displayed ligands and PEG coronas significantly impact targeting performance across multiple biological scales. Our work holds general implications for the rational design of receptor-targeted nanocarriers and also addresses challenges that are unique to the display of modular ligands anchored to vehicles bearing a common hydrophilic corona (here, PEG2k). We further demonstrate that the assessment of differential proteolysis kinetics provides a powerful tool for quantifying the relative biochemical access of targeting ligand prototypes displayed on nanocarrier surfaces. The biochemical access of a targeting ligand is an engineerable property that can be leveraged to control nanocarrier engagement with receptors on the target cell type over a continuous domain. When applied to the development of glaucoma nanotherapies using our FLT4-targeting approach, improving the biochemical access of peptide ligands increased drug delivery to Schlemm’s canal endothelial cells while minimizing off-target interactions with vascular endothelial cells. These performance improvements at the molecular and cellular levels translated to efficacy enhancements in a paired, longitudinal IOP study *in vivo*, thereby demonstrating the potential utility of our methods in clinically relevant settings. The technologies developed herein show promise for improving the potency and specificity of cell softening agents used to manage ocular hypertension. More generally, our rational design principles and methodology can be extended to the development of other targeted nanotherapies to solve diverse challenges in drug delivery.

## Supporting information

Vincent et al 2021 - Supplementary Information

## ASSOCIATED CONTENT

### Supporting Information

Supporting Information is available from the author.

Supplementary Methods for Dynamic Light Scattering (DLS) and Electrophoretic Light Scattering (ELS); Cryogenic Transmission Electron Microscopy (Cryo-TEM); Fourier Transform Infrared (FTIR) Spectroscopy; Animals; Ocular Histology Studies. Supplementary Figures including lipid-anchored FLT4-binding peptide constructs; Fourier-transform infrared (FTIR) spectroscopy spectra of purified micelle formulations; fluorescence emission scan of FLT4-binding peptide compared to blank PEG-*b*-PPS micelles; targeting peptide calibration; Small angle x-ray scattering (SAXS) of PEG-*b*-PPS micelles displaying FLT4-binding peptides at a 1% molar ratio; LatA concentration series for calibrating HPLC measurements; baseline intraocular pressure (IOP) comparison between left and right eyes of C57BL/6J mice. Supplementary Tables including PEG-*b*-PPS Micelle (MC) diblock copolymer; FLT4-targeting peptide constructs.

## Author Contributions

MPV prepared and characterized all nanocarrier formulations. MPV designed and conducted mass spectrometry-based peptide proteolysis assays for quantifying peptide accessibility. MPV conducted SAXS experiments at Argonne National Laboratory. WDS and KMP provided human Schlemm’s canal endothelial cells for use in this study. MPV conducted the FLT4 expression analysis. MPV, AV, and TS contributed to the primary cell culture *in vitro* studies. MPV and TS conducted nanocarrier uptake studies *in vitro*. MPV, MJ, and EAS contributed to study design. MPV, GL, WDS, RR, HG, MJ, and EAS contributed to the design and execution of the mouse studies. MPV and MJ contributed to the statistical analysis. MPV, MJ, EAS, WDS, GL, and HG contributed to writing of the manuscript. MPV created all figures and illustrations. All authors have given approval to the final version of the manuscript.

## Funding Sources

MPV gratefully acknowledges support from the Ryan Fellowship, the International Institute for Nanotechnology at Northwestern University, and the Northwestern University Multidisciplinary Visual Sciences Training Program (T32 Fellowship funded by NEI Award 2T32EY025202-06). This research was supported by the National Science Foundation CAREER Award no. 1453576 and the National Institutes of Health Director’s New Innovator Award no. 1DP2HL132390-01. Funding for in vitro studies was provided by NIH grant 1R01EY019696. Funding for animal studies was provided by NIH grant 1R0EY022359 and from BrightFocus Foundation grant G2019295. We thank Dr. Mark Karver (Northwestern University) for his assistance and support with peptide synthesis. We thank Eric W. Roth (Northwestern University) for his assistance with Cryo-TEM. Peptide synthesis was performed at the Peptide Synthesis Core Facility of the Simpson Querrey Institute at Northwestern University. This facility has current support from the Soft and Hybrid Nanotechnology Experimental (SHyNE) Resource (NSF ECCS-2025633). The Simpson Querrey Institute, Northwestern University Office for Research, U.S. Army Research Office, and the U.S. Army Medical Research and Materiel Command have also provided funding to develop this facility. This work made use of the EPIC facility of Northwestern University’s NUANCE Center, which has received support from the SHyNE Resource (NSF ECCS-2025633), the IIN, and Northwestern’s MRSEC program (NSF DMR-1720139). This work made use of the BioCryo facility of Northwestern University’s NUANCE Center, which has received support from the Soft and Hybrid Nanotechnology Experimental (SHyNE) Resource (NSF ECCS-1542205); the MRSEC program (NSF DMR-1720139) at the Materials Research Center; the International Institute for Nanotechnology (IIN); and the State of Illinois, through the IIN. This work made use of the IMSERC MS facility at Northwestern University, which has received support from the Soft and Hybrid Nanotechnology Experimental (SHyNE) Resource (NSF ECCS-2025633), the State of Illinois, and the International Institute for Nanotechnology (IIN). This work was further supported by the Northwestern University Robert H. Lurie Comprehensive Cancer Center (RHLCCC) Flow Cytometry Facility and a Cancer Center Support Grant (NCI CA060553). This work was further supported by the Northwestern University High Throughput Analysis Laboratory.

## Notes

The authors declare no conflict of interest.

## For Table of Contents Only

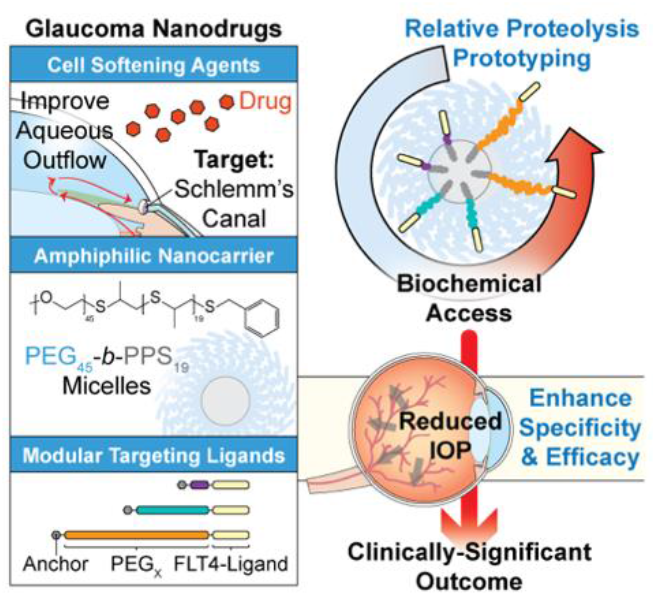

## Notes

### Competing Interest Statement

The authors have declared no competing interest.

## References

(1) Tham, Y.-C.; Li, X.; Wong, T. Y.; Quigley, H. A.; Aung, T.; Cheng, C.-Y. Global Prevalence of Glaucoma and Projections of Glaucoma Burden through 2040. Ophthalmology 2014, 121 (11), 2081–2090. https://doi.org/10.1016/j.ophtha.2014.05.013.

(2) Mao, L. K.; Stewart, W. C.; Shields, M. B. Correlation Between Intraocular Pressure Control and Progressive Glaucomatous Damage in Primary Open-Angle Glaucoma. Am J Ophthalmol 1991, 111 (1), 51–55. https://doi.org/10.1016/S0002-9394(14)76896-5.

(3) Quigley, H. A.; Nickells, R. W.; Kerrigan, L. A.; Pease, M. E.; Thibault, D. J.; Zack, D. J. Retinal Ganglion Cell Death in Experimental Glaucoma and after Axotomy Occurs by Apoptosis. Invest Ophthalmol Vis Sci 1995, 36 (5), 774–786.

(4) Garcia-Valenzuela, E.; Shareef, S.; Walsh, J.; Sharma, S. C. Programmed Cell Death of Retinal Ganglion Cells during Experimental Glaucoma. Exp Eye Res 1995, 61 (1), 33–44. https://doi.org/10.1016/S0014-4835(95)80056-5.

(5) Kerrigan, L. A.; Zack, D. J.; Quigley, H. A.; Smith, S. D.; Pease, M. E. TUNEL-Positive= Ganglion Cells in Human Primary Open-Angle Glaucoma. Arch Ophthalmol 1997, 115 (8), 1031–1035. https://doi.org/10.1001/archopht.1997.01100160201010.

(6) Guo, L.; Moss, S. E.; Alexander, R. A.; Ali, R. R.; Fitzke, F. W.; Cordeiro, M. F. Retinal Ganglion Cell Apoptosis in Glaucoma Is Related to Intraocular Pressure and IOP-Induced Effects on Extracellular Matrix. Invest Ophthalmol Vis Sci 2005, 46 (1), 175–182. https://doi.org/10.1167/iovs.04-0832.

(7) Allingham, R. R.; de Kater, A. W.; Ethier, C. R.; Anderson, P. J.; Hertzmark, E.; Epstein, D. L. The Relationship between Pore Density and Outflow Facility in Human Eyes. Invest Ophthalmol Vis Sci 1992, 33 (5), 1661–1669.

(8) Johnson, M.; Shapiro, A.; Ethier, C. R.; Kamm, R. D. Modulation of Outflow Resistance by the Pores of the Inner Wall Endothelium. Invest Ophthalmol Vis Sci 1992, 33 (5), 1670–1675.

(9) Sit, A. J.; Coloma, F. M.; Ethier, C. R.; Johnson, M. Factors Affecting the Pores of the Inner Wall Endothelium of Schlemm’s Canal. Invest Ophthalmol Vis Sci 1997, 38 (8), 1517–1525.

(10) Ethier, C. R.; Coloma, F. M.; Sit, A. J.; Johnson, M. Two Pore Types in the Inner-Wall Endothelium of Schlemm’s Canal. Invest Ophthalmol Vis Sci 1998, 39 (11), 2041–2048.

(11) Johnson, M.; Chan, D.; Read, A. T.; Christensen, C.; Sit, A.; Ethier, C. R. The Pore Density in the Inner Wall Endothelium of Schlemm’s Canal of Glaucomatous Eyes. Invest Ophthalmol Vis Sci 2002, 43 (9), 2950–2955.

(12) Johnson, M. “What Controls Aqueous Humour Outflow Resistance?”Exp Eye Res 2006, 82 (4), 545–557. https://doi.org/10.1016/j.exer.2005.10.011.

(13) Zhou, E. H.; Krishnan, R.; Stamer, W. D.; Perkumas, K. M.; Rajendran, K.; Nabhan, J. F.; Lu, Q.; Fredberg, J. J.; Johnson, M. Mechanical Responsiveness of the Endothelial Cell of Schlemm’s Canal: Scope, Variability and Its Potential Role in Controlling Aqueous Humour Outflow. J R Soc Interface 2012, 9 (71), 1144–1155. https://doi.org/10.1098/rsif.2011.0733.

(14) Braakman, S. T.; Pedrigi, R. M.; Read, A. T.; Smith, J. A. E.; Stamer, W. D.; Ethier, C. R.; Overby, D. R. Biomechanical Strain as a Trigger for Pore Formation in Schlemm’s Canal Endothelial Cells. Exp Eye Res 2014, 127, 224–235. https://doi.org/10.1016/j.exer.2014.08.003.

(15) Overby, D. R.; Zhou, E. H.; Vargas-Pinto, R.; Pedrigi, R. M.; Fuchshofer, R.; Braakman, S. T.; Gupta, R.; Perkumas, K. M.; Sherwood, J. M.; Vahabikashi, A.; Dang, Q.; Kim, J. H.; Ethier, C. R.; Stamer, W. D.; Fredberg, J. J.; Johnson, M. Altered Mechanobiology of Schlemm’s Canal Endothelial Cells in Glaucoma. Proc Natl Acad Sci U S A 2014, 111 (38), 13876–13881. https://doi.org/10.1073/pnas.1410602111.

(16) Vahabikashi, A.; Gelman, A.; Dong, B.; Gong, L.; Cha, E. D. K.; Schimmel, M.; Tamm, E. R.; Perkumas, K.; Stamer, W. D.; Sun, C.; Zhang, H. F.; Gong, H.; Johnson, M. Increased Stiffness and Flow Resistance of the Inner Wall of Schlemm’s Canal in Glaucomatous Human Eyes. Proc Natl Acad Sci U S A 2019, 116 (52), 26555–26563. https://doi.org/10.1073/pnas.1911837116.

(17) Schwalbe, G. Untersuchungen über die Lymphbahnen des Auges und ihre Begrenzungen. Archiv f. mikrosk. Anatomie 1870, 6 (1), 261–362. https://doi.org/10.1007/BF02955984.

(18) Leber, Th. Studien über den Flüssigkeitswechsel im Auge. Graefes Arhiv für Ophthalmologie 1873, 19 (2), 87–185. https://doi.org/10.1007/BF01720618.

(19) Seidel, E. Weitere experimentelle Untersuchungen über die Quelle und den Verlauf der intraokularen Saftströmung. Graefes Arhiv für Ophthalmologie 1921, 104 (2), 284–292. https://doi.org/10.1007/BF01858592.

(20) Ascher, K. W. Aqueous Veins. Am J Ophthalmol 1942, 25 (1), 31–38. https://doi.org/10.1016/S0002-9394(42)93294-X.

(21) Bill, A. The Albumin Exchange in the Rabbit Eye. Acta Physiol Scand 1964, 60 (1–2), 18–29. https://doi.org/10.1111/j.1748-1716.1964.tb02865.x.

(22) Bill, A. The Drainage of Albumin from the Uvea. Exp Eye Res 1964, 3 (2), 179–187. https://doi.org/10.1016/S0014-4835(64)80033-6.

(23) Bill, A. The Aqueous Humor Drainage Mechanism in the Cynomolgus Monkey (Macaca Irus) with Evidence for Unconventional Routes. Invest Ophthalmol 1965, 4 (5), 911–919.

(24) Bill, A.; Hellsing, K. Production and Drainage of Aqueous Humor in the Cynomolgus Monkey (Macaca Irus). Invest Ophthalmol 1965, 4 (5), 920–926.

(25) Grant, W. M. Clinical Measurements of Aqueous Outflow. AMA Arch Ophthalmol 1951, 46 (2), 113–131. https://doi.org/10.1001/archopht.1951.01700020119001.

(26) Tanna, A. P.; Johnson, M. Rho Kinase Inhibitors as a Novel Treatment for Glaucoma and Ocular Hypertension. Ophthalmology 2018, 125 (11), 1741–1756. https://doi.org/10.1016/j.ophtha.2018.04.040.

(27) Kaufman, P. L.; Bill, A.; Bárány, E. H. Effect of Cytochalasin B on Conventional Drainage of Aqueous Humor in the Cynomolgus Monkey. Exp. Eye Res. 1977, 25 Suppl, 411–414. https://doi.org/10.1016/s0014-4835(77)80037-7.

(28) Peterson, J. A.; Tian, B.; Bershadsky, A. D.; Volberg, T.; Gangnon, R. E.; Spector, I.; Geiger, B.; Kaufman, P. L. Latrunculin-A Increases Outflow Facility in the Monkey. Invest Ophthalmol Vis Sci 1999, 40 (5), 931–941.

(29) Peterson, J. A.; Tian, B.; Geiger, B.; Kaufman, P. L. Effect of Latrunculin-B on Outflow Facility in Monkeys. Exp Eye Res 2000, 70 (3), 307–313. https://doi.org/10.1006/exer.1999.0797.

(30) Levy, B.; Ramirez, N.; Novack, G. D.; Kopczynski, C. Ocular Hypotensive Safety and Systemic Absorption of AR-13324 Ophthalmic Solution in Normal Volunteers. Am J Ophthalmol 2015, 159 (5), 980–985.e1. https://doi.org/10.1016/j.ajo.2015.01.026.

(31) Bacharach, J.; Dubiner, H. B.; Levy, B.; Kopczynski, C. C.; Novack, G. D. Double-Masked, Randomized, Dose–Response Study of AR-13324 versus Latanoprost in Patients with Elevated Intraocular Pressure. Ophthalmology 2015, 122 (2), 302–307. https://doi.org/10.1016/j.ophtha.2014.08.022.

(32) Lewis, R. A.; Levy, B.; Ramirez, N.; Kopczynski, C. C.; Usner, D. W.; Novack, G. D.; Group, for the P.-C. S. Fixed-Dose Combination of AR-13324 and Latanoprost: A Double-Masked, 28-Day, Randomised, Controlled Study in Patients with Open-Angle Glaucoma or Ocular Hypertension. Br J Ophthalmol 2016, 100 (3), 339–344. https://doi.org/10.1136/bjophthalmol-2015-306778.

(33) Serle, J. B.; Katz, L. J.; McLaurin, E.; Heah, T.; Ramirez-Davis, N.; Usner, D. W.; Novack, G. D.; Kopczynski, C. C. Two Phase 3 Clinical Trials Comparing the Safety and Efficacy of Netarsudil to Timolol in Patients With Elevated Intraocular Pressure: Rho Kinase Elevated IOP Treatment Trial 1 and 2 (ROCKET-1 and ROCKET-2). Am J Ophthalmol 2018, 186, 116–127. https://doi.org/10.1016/j.ajo.2017.11.019.

(34) Kahook, M. Y.; Serle, J. B.; Mah, F. S.; Kim, T.; Raizman, M. B.; Heah, T.; Ramirez-Davis, N.; Kopczynski, C. C.; Usner, D. W.; Novack, G. D.; ROCKET-2 Study Group. Long-Term Safety and Ocular Hypotensive Efficacy Evaluation of Netarsudil Ophthalmic Solution: Rho Kinase Elevated IOP Treatment Trial (ROCKET-2). Am J Ophthalmol 2019, 200, 130–137. https://doi.org/10.1016/j.ajo.2019.01.003.

(35) Asrani, S.; Robin, A. L.; Serle, J. B.; Lewis, R. A.; Usner, D. W.; Kopczynski, C. C.; Heah, T.; Ackerman, S. L.; Alpern, L. M.; Asrani, S.; Bashford, K.; Bluestein, E. C.; Boyce, J. D.; Branch, J. D.; Brubaker, J. W.; Christie, W. C.; Cohen, J. S.; Collins, N. M.; Corin, S. M.; Daynes, T. E.; Depenbusch, M.; Dixon, E.-R.; Duzman, E.; Flowers, B. E.; Flynn, W. J.; Fong, R.; Gira, J. P.; Goldberg, D. F.; Greene, B.; Han, S. B.; Henderson, T. T.; Jerkins, G.; Jong, K. Y.; Katzen, L. B.; Khemsara, V.; Klugo, K. L.; Kozlovsky, J. F.; Leonardo, D.; Liu, Y.; LoBue, T. D.; Luchs, J. I.; Malhotra, R. P.; Mays, A.; McLaurin, E. B.; McMenemy, M. G.; Modi, S.; Moroi, S.; Mulaney, J.; Nagi, K.; Nicolau, J.; Parikh, M.; Patel, J. R.; Peplinski, L. S.; Perez, B. R.; Piltz-Seymour, J.; Sadri, E.; Saltzmann, R. M.; Schenker, H. I.; Swanic, M. J.; Tekwani, N.; Teymoorian, S.; Thomas, J. W.; Tyson, F. C.; Vold, S.; Weiss, M. J.; Zaman, F. Netarsudil/Latanoprost Fixed-Dose Combination for Elevated Intraocular Pressure: Three-Month Data from a Randomized Phase 3 Trial. Am J Ophthalmol 2019, 207, 248–257. https://doi.org/10.1016/j.ajo.2019.06.016.

(36) Tanihara, H.; Inoue, T.; Yamamoto, T.; Kuwayama, Y.; Abe, H.; Araie, M.; K-115 Clinical Study Group. Phase 2 Randomized Clinical Study of a Rho Kinase Inhibitor, K-115, in Primary Open-Angle Glaucoma and Ocular Hypertension. Am J Ophthalmol 2013, 156 (4), 731–736. https://doi.org/10.1016/j.ajo.2013.05.016.

(37) Stack, T.; Vincent, M.; Vahabikashi, A.; Li, G.; Perkumas, K. M.; Stamer, W. D.; Johnson, M.; Scott, E. Targeted Delivery of Cell Softening Micelles to Schlemm’s Canal Endothelial Cells for Treatment of Glaucoma. Small 2020, e2004205. https://doi.org/10.1002/smll.202004205.

(38) Aspelund, A.; Tammela, T.; Antila, S.; Nurmi, H.; Leppänen, V.-M.; Zarkada, G.; Stanczuk, L.; Francois, M.; Mäkinen, T.; Saharinen, P.; Immonen, I.; Alitalo, K. The Schlemm’s Canal Is a VEGF-C/VEGFR-3–Responsive Lymphatic-like Vessel. J Clin Invest 2014, 124 (9), 3975–3986. https://doi.org/10.1172/JCI75395.

(39) Patel, G.; Fury, W.; Yang, H.; Gomez-Caraballo, M.; Bai, Y.; Yang, T.; Adler, C.; Wei, Y.; Ni, M.; Schmitt, H.; Hu, Y.; Yancopoulos, G.; Stamer, W. D.; Romano, C. Molecular Taxonomy of Human Ocular Outflow Tissues Defined by Single-Cell Transcriptomics. Proc Natl Acad Sci U S A 2020, 117 (23), 12856–12867. https://doi.org/10.1073/pnas.2001896117.

(40) van Zyl, T.; Yan, W.; McAdams, A.; Peng, Y.-R.; Shekhar, K.; Regev, A.; Juric, D.; Sanes, J. R. Cell Atlas of Aqueous Humor Outflow Pathways in Eyes of Humans and Four Model Species Provides Insight into Glaucoma Pathogenesis. Proc Natl Acad Sci U S A 2020, 117 (19), 10339–10349. https://doi.org/10.1073/pnas.2001250117.

(41) Elias, D. R.; Poloukhtine, A.; Popik, V.; Tsourkas, A. Effect of Ligand Density, Receptor Density, and Nanoparticle Size on Cell Targeting. Nanomedicine 2013, 9 (2), 194–201. https://doi.org/10.1016/j.nano.2012.05.015.

(42) Huang, Y.; Jiang, K.; Zhang, X.; Chung, E. J. The Effect of Size, Charge, and Peptide Ligand Length on Kidney Targeting by Small, Organic Nanoparticles. Bioengineering & Transla Med 2020, 5 (3), e10173. https://doi.org/10.1002/btm2.10173.

(43) Boussommier-Calleja, A.; Bertrand, J.; Woodward, D. F.; Ethier, C. R.; Stamer, W. D.; Overby, D. R. Pharmacologic Manipulation of Conventional Outflow Facility in Ex Vivo Mouse Eyes. Invest Ophthalmol Vis Sci 2012, 53 (9), 5838–5845. https://doi.org/10.1167/iovs.12-9923.

(44) Boussommier-Calleja, A.; Li, G.; Wilson, A.; Ziskind, T.; Scinteie, O. E.; Ashpole, N. E.; Sherwood, J. M.; Farsiu, S.; Challa, P.; Gonzalez, P.; Downs, J. C.; Ethier, C. R.; Stamer, W. D.; Overby, D. R. Physical Factors Affecting Outflow Facility Measurements in Mice. Invest Ophthalmol Vis Sci 2015, 56 (13), 8331–8339. https://doi.org/10.1167/iovs.15-17106.

(45) Zhang, X.; Beckmann, L.; Miller, D. A.; Shao, G.; Cai, Z.; Sun, C.; Sheibani, N.; Liu, X.; Schuman, J.; Johnson, M.; Kume, T.; Zhang, H. F. In Vivo Imaging of Schlemm’s Canal and Limbal Vascular Network in Mouse Using Visible-Light OCT. Invest Ophthalmol Vis Sci 2020, 61 (2), 23–23. https://doi.org/10.1167/iovs.61.2.23.

(46) Scott, E. A.; Stano, A.; Gillard, M.; Maio-Liu, A. C.; Swartz, M. A.; Hubbell, J. A. Dendritic Cell Activation and T Cell Priming with Adjuvant- and Antigen-Loaded Oxidation-Sensitive Polymersomes. Biomaterials 2012, 33 (26), 6211–6219. https://doi.org/10.1016/j.biomaterials.2012.04.060.

(47) Yi, S.; Allen, S. D.; Liu, Y.-G.; Ouyang, B. Z.; Li, X.; Augsornworawat, P.; Thorp, E. B.; Scott, E. A. Tailoring Nanostructure Morphology for Enhanced Targeting of Dendritic Cells in Atherosclerosis. ACS Nano 2016, 10 (12), 11290–11303. https://doi.org/10.1021/acsnano.6b06451.

(48) Stack, T.; Vahabikashi, A.; Johnson, M.; Scott, E. Modulation of Schlemm’s Canal Endothelial Cell Stiffness via Latrunculin Loaded Block Copolymer Micelles. J Biomed Mater Res A 2018, 106 (7), 1771–1779. https://doi.org/10.1002/jbm.a.36376.

(49) Stamer, W. D.; Roberts, B. C.; Howell, D. N.; Epstein, D. L. Isolation, Culture, and Characterization of Endothelial Cells from Schlemm’s Canal. Invest Ophthalmol Vis Sci 1998, 39 (10), 1804–1812.

(50) Perkumas, K. M.; Stamer, W. D. Protein Markers and Differentiation in Culture for Schlemm’s Canal Endothelial Cells. Exp Eye Res 2012, 96 (1), 82–87. https://doi.org/10.1016/j.exer.2011.12.017.

(51) Vincent, M. P.; Bobbala, S.; Karabin, N. B.; Frey, M.; Liu, Y.; Navidzadeh, J. O.; Stack, T.; Scott, E. A. Surface Chemistry-Mediated Modulation of Adsorbed Albumin Folding State Specifies Nanocarrier Clearance by Distinct Macrophage Subsets. Nat Commun 2021, 12 (1), 648. https://doi.org/10.1038/s41467-020-20886-7.

(52) Chen, T. J.; Kotecha, N. Cytobank: Providing an Analytics Platform for Community Cytometry Data Analysis and Collaboration. Curr Top Microbiol Immunol 2014, 377, 127–157. https://doi.org/10.1007/82_2014_364.

(53) Kizhatil, K.; Ryan, M.; Marchant, J. K.; Henrich, S.; John, S. W. M. Schlemm’s Canal Is a Unique Vessel with a Combination of Blood Vascular and Lymphatic Phenotypes That Forms by a Novel Developmental Process. PLoS Biol 2014, 12 (7). https://doi.org/10.1371/journal.pbio.1001912.

(54) Kashman, Y.; Groweiss, A.; Shmueli, U. Latrunculin, a New 2-Thiazolidinone Macrolide from the Marine Sponge Latrunculia Magnifica. Tetrahedron Lett 1980, 21(37), 3629–3632. https://doi.org/10.1016/0040-4039(80)80255-3.

(55) Spector, I.; Shochet, N. R.; Kashman, Y.; Groweiss, A. Latrunculins: Novel Marine Toxins That Disrupt Microfilament Organization in Cultured Cells. Science 1983, 219 (4584), 493–495. https://doi.org/10.1126/science.6681676.

(56) Coué, M.; Brenner, S. L.; Spector, I.; Korn, E. D. Inhibition of Actin Polymerization by Latrunculin A. FEBS Letters 1987, 213 (2), 316–318. https://doi.org/10.1016/0014-5793(87)81513-2.

(57) Yarmola, E. G.; Somasundaram, T.; Boring, T. A.; Spector, I.; Bubb, M. R. Actin-Latrunculin A Structure and Function. Differential Modulation of Actin-Binding Protein Function by Latrunculin A. J Biol Chem 2000, 275 (36), 28120–28127. https://doi.org/10.1074/jbc.M004253200.

(58) Petros, R. A.; DeSimone, J. M. Strategies in the Design of Nanoparticles for Therapeutic Applications. Nat Rev Drug Discov 2010, 9 (8), 615–627. https://doi.org/10.1038/nrd2591.

(59) Yi, S.; Zhang, X.; Sangji, M. H.; Liu, Y.; Allen, S. D.; Xiao, B.; Bobbala, S.; Braverman, C. L.; Cai, L.; Hecker, P. I.; DeBerge, M.; Thorp, E. B.; Temel, R. E.; Stupp, S. I.; Scott, E. A. Surface Engineered Polymersomes for Enhanced Modulation of Dendritic Cells During Cardiovascular Immunotherapy. Adv Funct Mater 2019, 29(42), 1904399. https://doi.org/10.1002/adfm.201904399.

(60) Allen, S.; Vincent, M.; Scott, E. Rapid, Scalable Assembly and Loading of Bioactive Proteins and Immunostimulants into Diverse Synthetic Nanocarriers Via Flash Nanoprecipitation. J Vis Exp 2018, No. 138, e57793. https://doi.org/10.3791/57793.

(61) Park, C. Y.; Zhou, E. H.; Tambe, D.; Chen, B.; Lavoie, T.; Dowell, M.; Simeonov, A.; Maloney, D. J.; Marinkovic, A.; Tschumperlin, D. J.; Burger, S.; Frykenberg, M.; Butler, J. P.; Stamer, W. D.; Johnson, M.; Solway, J.; Fredberg, J. J.; Krishnan, R. High-Throughput Screening for Modulators of Cellular Contractile Force. Integr Biol (Camb) 2015, 7 (10), 1318–1324. https://doi.org/10.1039/c5ib00054h.

(62) Hämäläinen, K. M.; Kananen, K.; Auriola, S.; Kontturi, K.; Urtti, A. Characterization of Paracellular and Aqueous Penetration Routes in Cornea, Conjunctiva, and Sclera. Invest Ophthalmol Vis Sci 1997, 38 (3), 627–634.

(63) Prausnitz, M. R.; Noonan, J. S. Permeability of Cornea, Sclera, and Conjunctiva: A Literature Analysis for Drug Delivery to the Eye. J Pharm Sci 1998, 87 (12), 1479–1488. https://doi.org/10.1021/js9802594.

(64) Agrahari, V.; Mandal, A.; Agrahari, V.; Trinh, H. M.; Joseph, M.; Ray, A.; Hadji, H.; Mitra, R.; Pal, D.; Mitra, A. K. A Comprehensive Insight on Ocular Pharmacokinetics. Drug Deliv Transl Res 2016, 6 (6), 735–754. https://doi.org/10.1007/s13346-016-0339-2.

(65) Velluto, D.; Thomas, S. N.; Simeoni, E.; Swartz, M. A.; Hubbell, J. A. PEG-b-PPS-b-PEI Micelles and PEG-b-PPS/PEG-b-PPS-b-PEI Mixed Micelles as Non-Viral Vectors for Plasmid DNA: Tumor Immunotoxicity in B16F10 Melanoma. Biomaterials 2011, 32 (36), 9839–9847. https://doi.org/10.1016/j.biomaterials.2011.08.079.

(66) Wilson, A. M.; Di Polo, A. Gene Therapy for Retinal Ganglion Cell Neuroprotection in Glaucoma. Gene Ther 2012, 19 (2), 127–136. https://doi.org/10.1038/gt.2011.142.

(67) Choquet, H.; Wiggs, J. L.; Khawaja, A. P. Clinical Implications of Recent Advances in Primary Open-Angle Glaucoma Genetics. Eye 2020, 34 (1), 29–39. https://doi.org/10.1038/s41433-019-0632-7.

(68) Napoli, A.; Valentini, M.; Tirelli, N.; Müller, M.; Hubbell, J. A. Oxidation-Responsive Polymeric Vesicles. Nature Materials 2004, 3 (3), 183–189. https://doi.org/10.1038/nmat1081.

(69) Vasdekis, A. E.; Scott, E. A.; O’Neil, C. P.; Psaltis, D.; Hubbell, Jeffrey. A. Precision Intracellular Delivery Based on Optofluidic Polymersome Rupture. ACS Nano 2012, 6 (9), 7850–7857. https://doi.org/10.1021/nn302122h.

(70) Bobbala, S.; Allen, S. D.; Scott, E. A. Flash Nanoprecipitation Permits Versatile Assembly and Loading of Polymeric Bicontinuous Cubic Nanospheres. Nanoscale 2018, 10 (11), 5078–5088. https://doi.org/10.1039/C7NR06779H.

(71) Allen, S. D.; Bobbala, S.; Karabin, N. B.; Modak, M.; Scott, E. A. Benchmarking Bicontinuous Nanospheres against Polymersomes for in Vivo Biodistribution and Dual Intracellular Delivery of Lipophilic and Water-Soluble Payloads. ACS Appl Mater Interfaces 2018, 10 (40), 33857–33866. https://doi.org/10.1021/acsami.8b09906.

(72) Bobbala, S.; Allen, S. D.; Yi, S.; Vincent, M.; Frey, M.; Karabin, N. B.; Scott, E. A. Employing Bicontinuous-to-Micellar Transitions in Nanostructure Morphology for on-Demand Photo-Oxidation Responsive Cytosolic Delivery and off–on Cytotoxicity. Nanoscale 2020, 12 (9), 5332–5340. https://doi.org/10.1039/C9NR10921H.

