## Supplementary material for "Surface Engineering of FLT4-Targeted Nanocarriers Enhances Cell-Softening Glaucoma Therapy": Vincent et al 2021 - Supplementary Information

**Supplementary Methods**

**Dynamic Light Scattering (DLS) and Electrophoretic Light Scattering (ELS).** The number-average diameter and polydispersity index (PDI) of nanocarriers was determined by DLS using a Zetasizer Nano instrument (Malvern Instruments). The number average PDI was calculated using a custom matlab script (version R2019a). ELS was performed to determine the zeta potential of nanocarriers diluted (1:10) in 0.1x phosphate buffered saline (PBS).

**Cryogenic Transmission Electron Microscopy (Cryo-TEM).** Copper grids (200 mesh) with a lacey carbon membrane (EMS Cat# LC200-Cu-100) were glow-discharged in a Pelco easiGlow glow discharger (Ted Pella). This procedure used an atmosphere plasma generated at 15 mA for 15 seconds and a pressure of 0.24 mbar. A sample volume of 4 µL was applied to the grid. The grid was blotted for 5 s with a blot offset of +0.5 mm and was subsequently plunged into liquid ethane within a FEI Vitrobot Mark III plunge freezing instrument (Thermo Fisher Scientific). Specimens were imaged using a JEOL JEM1230 LaB6 emission TEM (JEOL USA) operating at 100 keV. The plunge-frozen grids were kept vitreous at -180 °C in a Gatan Cryo Transfer Holder model 626.6 (Gatan) during imaging. Images were acquired using a Gatan Orius SC1000 CCD camera Model 831 (Gatan).

**Fourier Transform Infrared (FTIR) Spectroscopy.** FTIR spectroscopy was performed using a Nicolet iS50 FTIR Spectrometer (Thermo Scientific). The background spectra for air was collected prior to obtaining measurements for the experimental samples. Spectra were obtained from liquid samples of purified PEG-*b*-PPS micelle formulations prepared with and without the specified targeting peptide construct (PG6, PG24, or PG48) displayed at a 5% molar ratio (peptide:polymer). Measurements were obtained in the 2000-600 $cm^{-1}$ range and 64 scans were collected per sample. The presence of peptide bonds was detected by analyzing the amide I (1700-1600 $cm^{-1}$) and amide II (1590-1520 $cm^{-1}$) bands.

**Animals.** All experiments were completed in compliance with the Association for Research in Vision and Ophthalmology (ARVO) Statement for the Use of Animals in

Ophthalmic and Vision Research. For IOP studies, mice were handled in accordance with an approved protocol (A001-19-01) by Institutional Animal Care and Use Committee of Duke University. C57BL/6 (C57) mice were purchased from The Jackson Laboratory. Animals were housed and bred in clear cages, and were maintained in housing rooms at 21 °C with a 12 h:12 h light:dark cycle.

For ocular histology studies, local Institutional Animal Care and Use Committee (IACUC) approval was obtained. Two C57BL/6 mice (7-week-old) were purchased from The Jackson Laboratory (Bar Harbor, ME). Mice were housed in the Animal Science Center of Boston University Medical Campus, with a 12-hour light/12-hour dark cycle and access to food and water ad libitum. Upon arrival, mice were examined to confirm a normal appearance, i.e. free of any signs of ocular disease, and allowed to acclimatize for at least three days before experiments.

**Ocular Histology Studies.** Mice (n=2) were anesthetized with 87.5 mg/kg ketamine (Henry Schein Inc.) and 12.5 mg/kg xylazine (Henry Schein Inc.) by intraperitoneal injection. One eye of each mouse was then injected with nanocarriers by adapting previously published procedure for fluorescent tracer injection[1]. In brief, a 10-µl Hamilton microsyringe (Nanofil; World Precision Instruments) was loaded with 1 µl solution of nanocarriers (20mg/ml), as well as 2 µl modified 4% paraformaldehyde (PFA) separated by a 0.2 µl air bubble. A 35G needle (NF35BL- 2; World Precision Instruments) connected to this syringe was then inserted into the anterior chamber centrally to optimize uniform distribution. The nanoparticle volume was delivered at 4 nl/s by a microprocessor-based microsyringe pump controller (Micro4; World Precision Instruments). The 12 o'clock position of the eye was marked using TMD Tissue Marking Dye (Triangle Biomedical Sciences) to provide orientation. 45 minutes were allowed for the nanocarriers to migrate through the anterior chamber, penetrate the trabecular meshwork, and reach Schlemm's canal while the needle remained in the eye. During this time, artificial tears (Henry Schein Inc.) were applied to the cornea of both eyes to prevent dehydration. 4%

PFA (2 μl) was subsequently injected into the anterior chamber while additional fixative was simultaneously applied to the exterior of the eye. After 30 minutes of fixation, both eyes were enucleated after euthanization of the mouse, immersion-fixed in 4% PFA at 4 °C overnight, then transferred in PBS and kept at 4 °C for further processing.

### Supplementary Figures

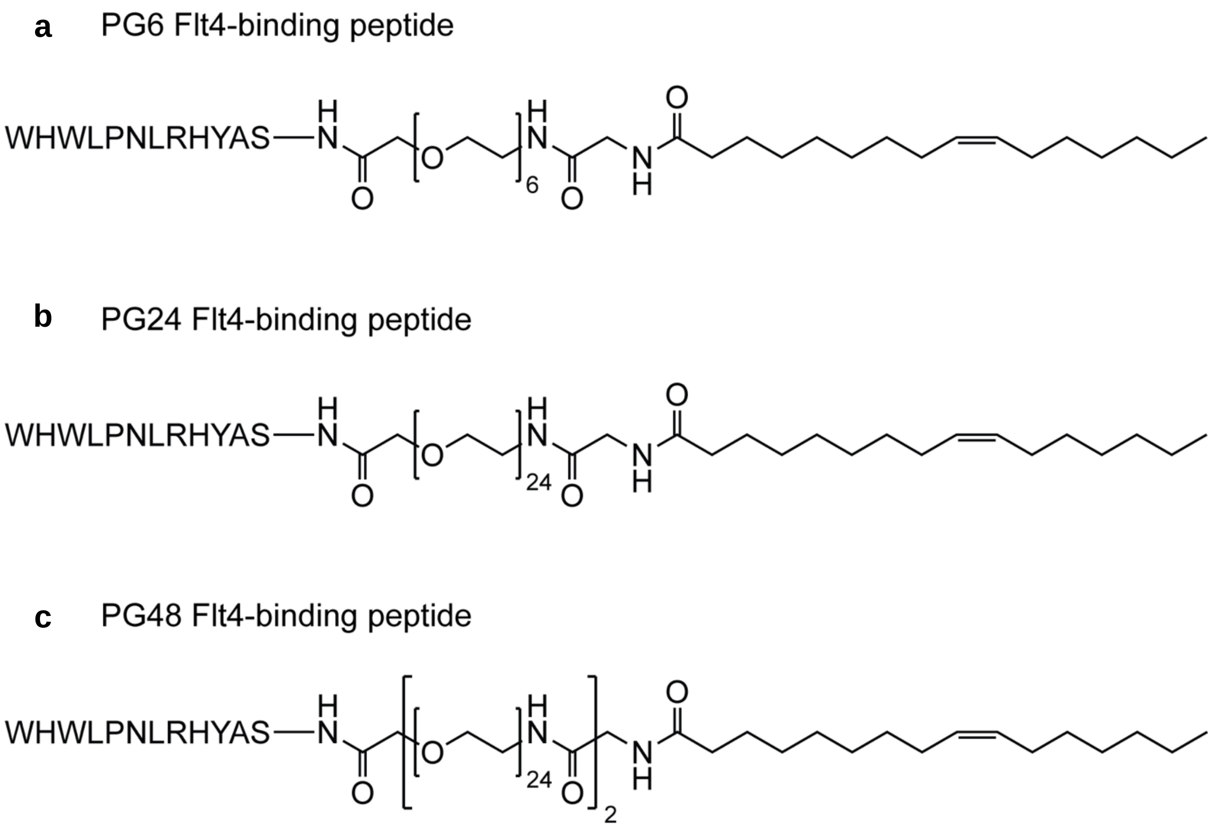


**Figure S1. Lipid-anchored FLT4-binding peptide constructs.** Lipid anchored peptide constructs were synthesized of the form [palmitoleic acid]-[$PEG_x$ spacer]-[Flt4-binding peptide] (listed N-terminus to C-terminus). The constructs differed by the length of their PEG spacers. (**a**) “PG6” Flt4-binding peptide (x = 6 PEG units), (**b**) “PG24” Flt4-binding peptide (x = 24 PEG units), and (**c**) “PG48” Flt4-binding peptide (x = 24x2 PEG units).

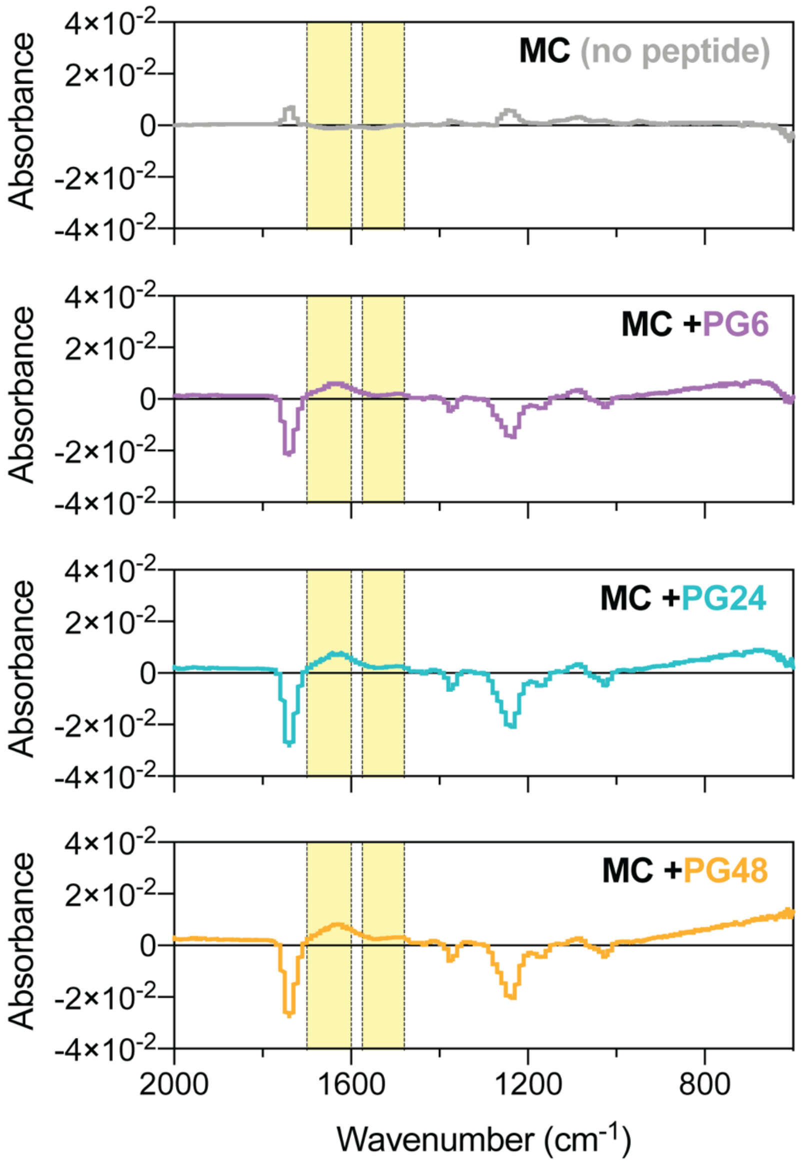


**Figure S2. Fourier-transform infrared (FTIR) spectroscopy spectra of purified micelle formulations.** The FTIR spectra of phosphate buffered saline (i.e. background from the buffer) was subtracted from the spectra of each of the specified formulations. In each case, the average spectrum is presented from a total of 64 scans. The amide I (1700-1600 cm$^{-1}$) and amide II (1590-1520 cm$^{-1}$) bands are highlighted in yellow.

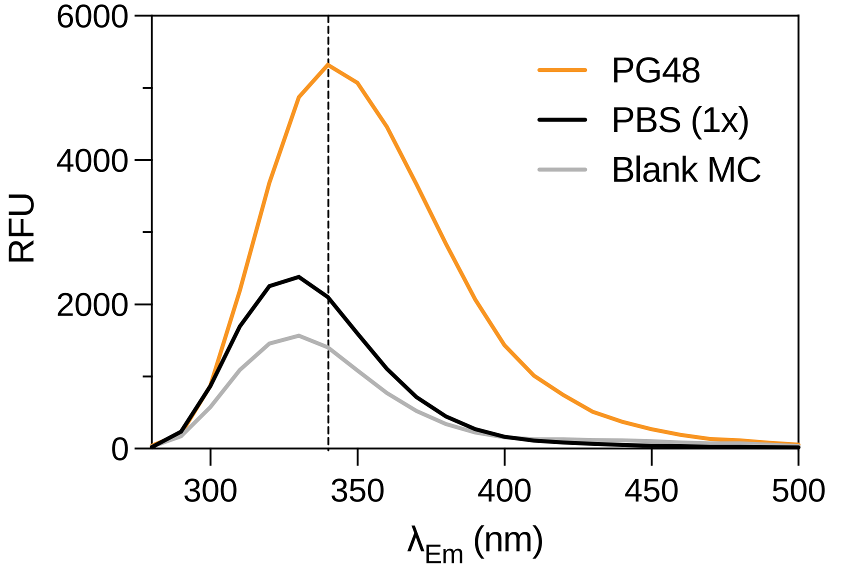


**Figure S3. Fluorescence emission scan of FLT4-binding peptide compared to blank PEG-*b*-PPS micelles.** The relative fluorescence signal is displayed for PBS (1x) background, blank micelles (MC), and the PG48 peptide. Fixed excitation wavelength ($\lambda_{Ex}$) = 270 nm. Emission scan: $\lambda_{Em}$ = 280 – 500 nm.

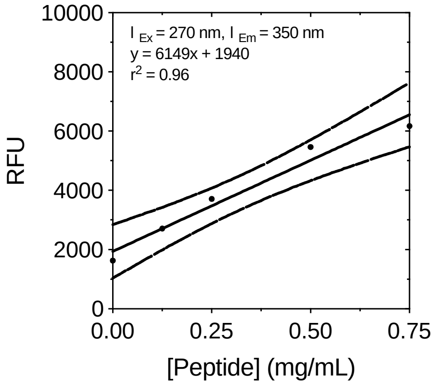


**Figure S4. Targeting peptide calibration.** A representative calibration curve is displayed for a peptide concentration series (0, 0.125, 0.25, 0.50, and 0.75 mg/mL). Data was acquired at an excitation wavelength ($\lambda_{Ex}$) of 270 nm and an emission wavelength ($\lambda_{Em}$) of 350 nm. The mean ± s.e.m. (n=3) is displayed with the 95% confidence bands. Where error bars are not displayed, the error is so small that the bars would not be visible. A linear regression model was fit to the data.

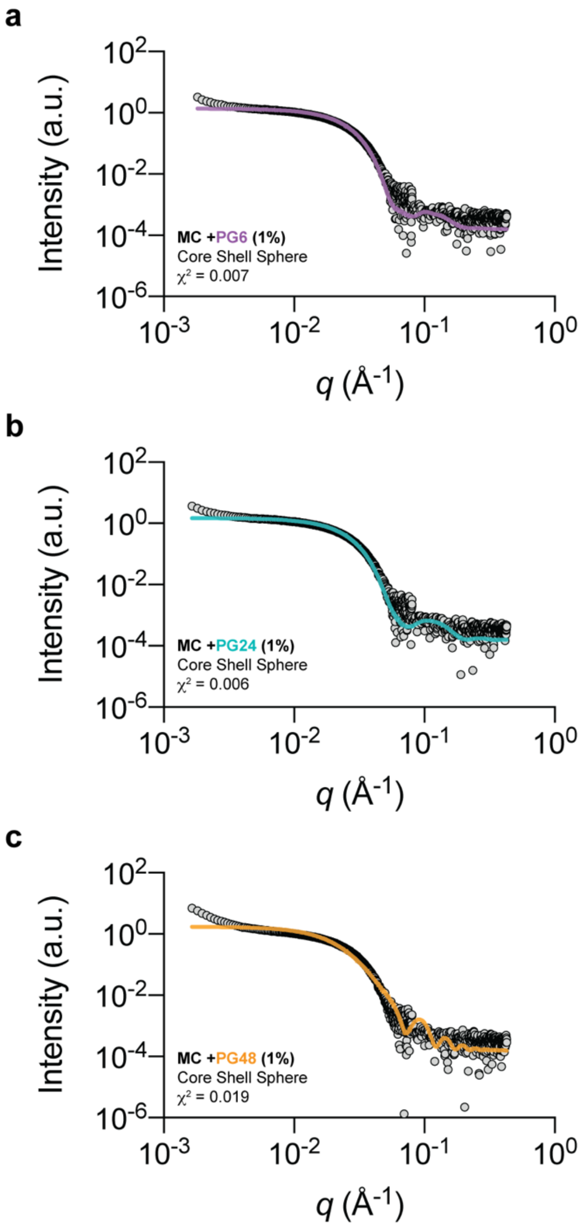


**Figure S5. Small angle x-ray scattering (SAXS) of PEG-*b*-PPS micelles displaying FLT4-binding peptides at a 1% molar ratio.** Core shell sphere models were fit to SAXS data obtained for PEG-b-PPS MCs displaying (**a**) PG6, (**b**) PG24, or (**c**) PG48 at a 1% molar ratio (peptide:polymer). The chi square of each model fit is inset ($\chi^2$ < 1.0 indicates a good model fit). See **Table 2** for more details regarding each model fit. SAXS was performed using synchrotron radiation at Argonne National Laboratory.

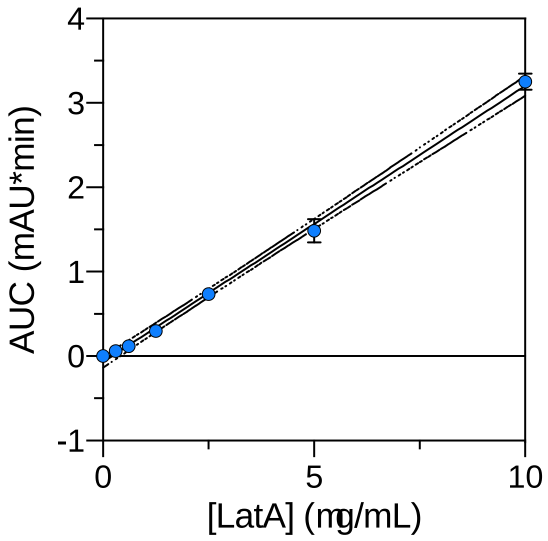


**Figure S6. LatA concentration series for calibrating HPLC measurements.** A representative calibration curve is displayed for a LatA concentration series prepared in methanol (0.0, 0.306, 0.612, 1.25, 2.5, 5.0, 10.0 μg/mL). The area under the curve (AUC) was obtained from three replicates (n=3). The mean ± s.e.m. is displayed with the 95% confidence bands. Where error bars are not displayed, the error is so small that the bars would not be visible. A linear regression model was fit to the data: y = 0.3273x – 0.06997, $r^2$ = 0.99.

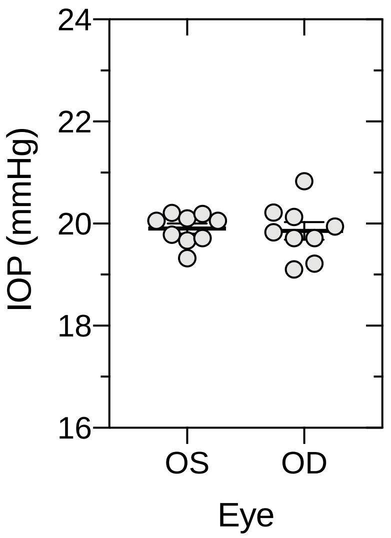


**Figure S7. Baseline intraocular pressure (IOP) comparison between left and right eyes of C57BL/6J mice.** Significant differences in the IOP of the left (OS) and right (OD) eyes of the animals were determined at baseline using a paired t-test and a 5% significance level (n=9). Significant differences in baseline IOP were not found.

### Supplementary Tables

**Table S1. PEG-*b*-PPS Micelle (MC) diblock copolymer.**

| BCP | $f_{PEG}$† | MW (g/mol) |
|---|---|---|
| MeO-$PEG_{45}$-*b*-$PPS_{19}$-Bn | 0.59 | 3,596 |

†Hydrophilic weight fraction of polymer.

**Table S2. FLT4-targeting peptide constructs.**

| Construct† | Lipid Anchor†† | PEG Spacer††† | | Peptide primary structure‡‡ | MW (g/mol) | Purity* |
|---|---|---|---|---|---|---|
| | | Units | Length (Å) | | | |
| PG6 | PA | 6 | 25.046 | WHWLPNLRHYAS | 2151.1 | >95% |
| PG24 | PA | 24 | 88.913 | WHWLPNLRHYAS | 3046.7 | >95% |
| PG48 | PA | 48‡ | 174.058 | WHWLPNLRHYAS | 4071.4 | >95% |

†All peptide constructs are of the form [Lipid]-[$PEG_X$ spacer]-[Peptide].
††Palmitoleic acid (PA).
†††The spacer length is presented as the number of PEG units.
‡The peptide construct with a 48-unit PEG spacer was synthesized using Fmoc-PEG24x2.
‡‡Peptide primary structure is listed N-terminus to C-terminus.
*Peptide purity determined using LC-MS.

**Supplementary References**

(1) Swaminathan, S. S.; Oh, D.-J.; Kang, M. H.; Ren, R.; Jin, R.; Gong, H.; Rhee, D. J. Secreted Protein Acidic and Rich in Cysteine (SPARC)-Null Mice Exhibit More Uniform Outflow. *Invest Ophthalmol Vis Sci* **2013**, *54* (3), 2035–2047. https://doi.org/10.1167/iovs.12-10950.
